# Destabilized host-parasite dynamics in newly founded populations

**DOI:** 10.1101/2024.06.24.600494

**Authors:** Daniel I. Bolnick, Rowan D.H. Barrett, Emma Choi, Lucas Eckert, Andrew P. Hendry, Emily V. Kerns, Åsa J. Lind, Kathryn Milligan-McClellan, Catherine L. Peichel, Kristofer Sasser, Alice R Thornton, Cole Wolf, Natalie C. Steinel, Jesse N. Weber

**Affiliations:** Department of Ecology and Evolutionary Biology, University of Connecticut Storrs CT, USA; Department of Biology, McGill University, Montreal, Quebec, Canada; Department of Integrative Biology, University of Wisconsin, Madison, WI, USA; Division of Evolutionary Ecology, Institute of Ecology and Evolution, University of Bern, Bern, Switzerland; Department of Molecular and Cell Biology, University of Connecticut, Storrs, CT, USA; Center for Pathogen Research and Training, University of Massachusetts, Lowell, MA, USA; Department of Biology, University of Massachusetts, Lowell, USA

## Abstract

When species disperse into previously unoccupied habitats, new populations encounter unfamiliar species interactions such as altered parasite loads. Theory predicts that newly founded populations should exhibit destabilized eco-evolutionary fluctuations in infection rates and immune traits. However, to understand founder effects biologists typically rely on retrospective studies of range expansions, missing early-generation infection dynamics. To remedy this, we experimentally founded whole-lake populations of threespine stickleback. Infection rates were temporally stable in native source lakes. In contrast, newly founded populations exhibit destabilized host-parasite dynamics: high starting infection rates led to increases in a heritable immune trait (peritoneal fibrosis), suppressing infection rates. The resulting temporal auto-correlation between infection and immunity suggest that newly founded populations can exhibit rapid host-parasite eco-evolutionary dynamics.

## Main Text

Species frequently disperse into unoccupied habitats, founding new populations that enable metapopulation persistence, geographic range expansion, invasive species establishment, and can be leveraged to reintroduce locally extinct species for conservation (*1*, *2*). Newly founded populations may be poorly adapted to their novel ecological conditions. In particular, immigrants experience altered parasite communities (*3*) which impose strong selection on host immunity (*4*, *5*). Eco-evolutionary models suggest that new host populations may rapidly evolve resistance to their new parasite community, which in turn should change parasite abundance. Consequently new populations may exhibit transiently destabilized host-parasite dynamics (*6–8*). Testing these predictions requires large-scale experiments that create new host populations and track subsequent changes in infection and immunity. We initiated such an experiment, reintroducing native threespine stickleback fish (*Gasterosteus aculeatus*) into eight recently-fishless lakes (Fig. 1A). Here, we show that founding new populations resulted in destabilized parasite dynamics and evolution of a heritable immune trait, in accord with eco-evolutionary theory.

**Fig. 1.**
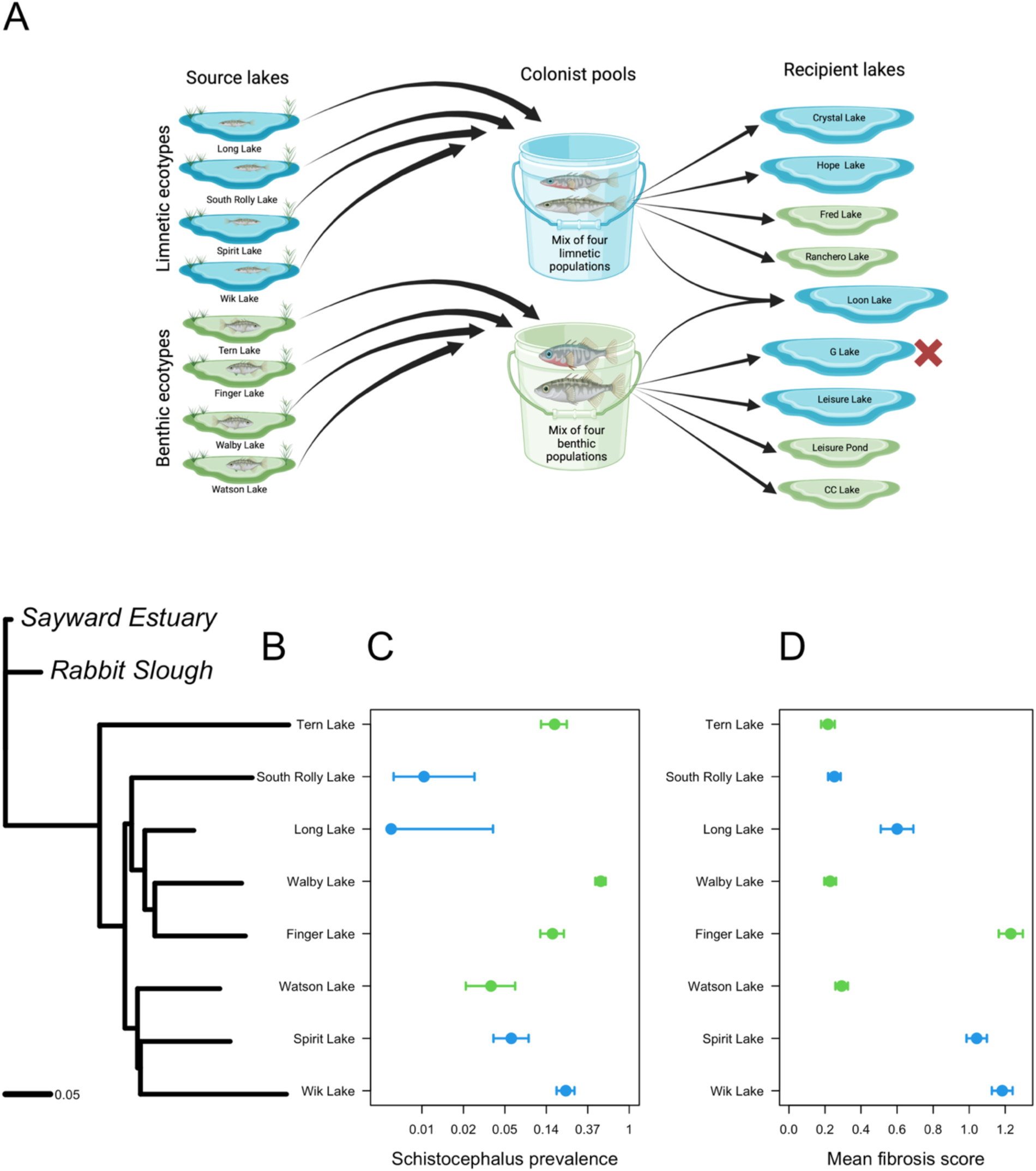
The experimental design and information about genetic, ecological, and immunological differences among the source lakes. (**A**) We identified eight lakes with native stickleback (map in fig. S1), which spanned an ecomorphological continuum from benthic to limnetic populations. In 2019 we collected and pooled equal numbers of fish from four benthic populations (and four limnetic populations) and factorially introduced these into four smaller and four larger recipient lakes (a ninth lake received both pools). The G Lake introduction failed. Created with BioRender.com. (**B)** A neighbor joining phylogenetic tree from PoolSeq data (*14*) showing genetic divergence between source lake populations, rooted by two marine anadromous populations from British Columbia (Sayward Estuary) and Alaska (Rabbit Slough). **(C)** Source populations differed in *Schistocephalus solidus* prevalence, ranging from 0% to 50% prevalence depending on the lake (binomial GLM lake effect Deviance = 378, df = 8, P < 0.0001). The x axis is on a log scale. **(D)** Fibrosis severity also differed among lakes, ranging from an average of 0.21 to 1.23 (Kruskal-Wallis c^2^ = 443, df = 8, P < 0.0001). We plot the means over four sample years (2019, 2021, 2022, 2023), with standard error bars, colored by population ecotype (green for limnetic, blue for benthic). Infection rates and fibrosis are not correlated with each other, or significantly associated with the morphologically-defined benthic/limnetic categorization (all P>0.1).

Threespine stickleback are hosts to diverse parasites communities (*9*), including a diphyllobothrian cestode, *Schistocephalus solidus*, which can grow to >50% of its host’s mass (*10*) and siphon >46% of the host’s baseline metabolic output (*11*). *S.solidus* infects stickleback when the fish ingests an infected copepod; the cestode penetrates the fish’s intestinal wall to grow in the body cavity. *S.solidus* reduces stickleback survival, growth, and fecundity (*12*). Consequently, freshwater stickleback populations repeatedly evolved increased resistance to *S.solidus* (*13*), including peritoneal fibrosis that reduces tapeworm growth and viability (*14*). This protective fibrosis is costly and irreversible (*15–17*), so some stickleback populations evolved tolerance instead, suppressing fibrosis and allowing tapeworm growth. As a result, stickleback exhibit heritable among-population differences in infection rate and fibrosis risk, providing a valuable model for understanding human fibrotic diseases (*18*). The eco-evolutionary reasons for these alternative fibrosis phenotypes in stickleback remain uncertain. A recent model suggests that when animal populations adapt to feed on locally available prey, their changing diet can alter parasite exposure rates, which in turn selects for immunity to commonly-encountered parasites (*19*). Stickleback populations inhabiting larger lakes tend to evolve a ‘limnetic’ body shape adapted to eating zooplankton (*20*, *21*), which include *S.solidus’* primary host copepods. Living in a limnetic habitat should increase tapeworm exposure risk (*22*), favoring evolutionary gain of fibrosis, thereby decreasing infection rates. The benthic ecotype, conversely, consumes macroinvertebrates in shallow-water habitats. With lower dietary exposure, benthic fish may evolve lower fibrosis, leaving them vulnerable when they do ingest a tapeworm. Thus, sticklebacks’ feeding ecology may contribute to the evolution of alternative fibrosis strategies. Newly founded populations may contain novel mixtures of immune and ecological genotypes which will not be precisely adapted to their novel habitat. Such populations should subsequently adapt to eating locally available prey, changing parasite exposure risks and driving evolution of immunity, which modifies infection rates (*19*).

To test the theory that new populations undergo destabilized host-parasite eco-evolutionary dynamics, in 2019 we reintroduced stickleback into newly fishless lakes on the Kenai Peninsula of Alaska (*23*), transplanting 10,831 fish from intact populations nearby (see Methods; Fig. 1A& S1). The eight source populations are genetically divergent (Fig. 1B), vary along a benthic-limnetic ecomorphological continuum (*24*), and differ in *S. solidus* prevalence (Fig. 1C) and fibrosis intensity (Fig. 1D). Infection rates were not significantly different between ecotypes (mean prevalence 0.17 and 0.03 in benthic and limnetic lakes respectively, t=1.15, P=0.323), despite limnetics’ greater exposure risk. This countergradient trend has been reported before (*25*), and fits a model in which high limnetic exposure risks drives evolution of greater immunity, negating or reversing the relationship between diet and successful infection (*19*). We created two mixed-population pools of founding stickleback, one pool drawn in approximately equal numbers from four benthic-ecotype lakes, the second pool drawn from four limnetic-ecotype lakes. These pools were introduced into eight fishless lakes, creating factorial combinations of benthic or limnetic pool fish added to small or large lakes (Fig. 1A, (*23*)). A ninth lake received both benthic and limnetic fish. We sampled source and recipient lake populations annually thereafter to track changes in both infection rates and fibrosis across four generations.

Cestode exposure induces fibrosis in some stickleback genotypes, but not others (*14*, *26*). In our source lakes, individual fish infected by tapeworms on average have 1.98-fold stronger fibrosis than those without (Fig. 2A), but this effect varied among lakes (1.1 to 8.2-fold). Consistent with our ecological predictions, fibrosis is higher in the larger lakes (Wik, Finger, and Spirit Lakes, Fig. 2B) where cestode exposure should be higher. By exposing lab-raised stickleback to *S.solidus*, we confirmed that fish from larger source lakes have a stronger fibrosis response to cestode exposure (Fig. 2C), whereas all populations responded similarly to a non-specific adjuvant (alum) that induces fibrosis (Figs. 2D). We conclude that there are heritable differences in severity of fibrosis response, and this variation is specific to *S. solidus.* Each pool of founding fish thus harbored genetic variation in fibrosis, enabling eco-evolutionary dynamics in the recipient lakes.

**Figure 2.**
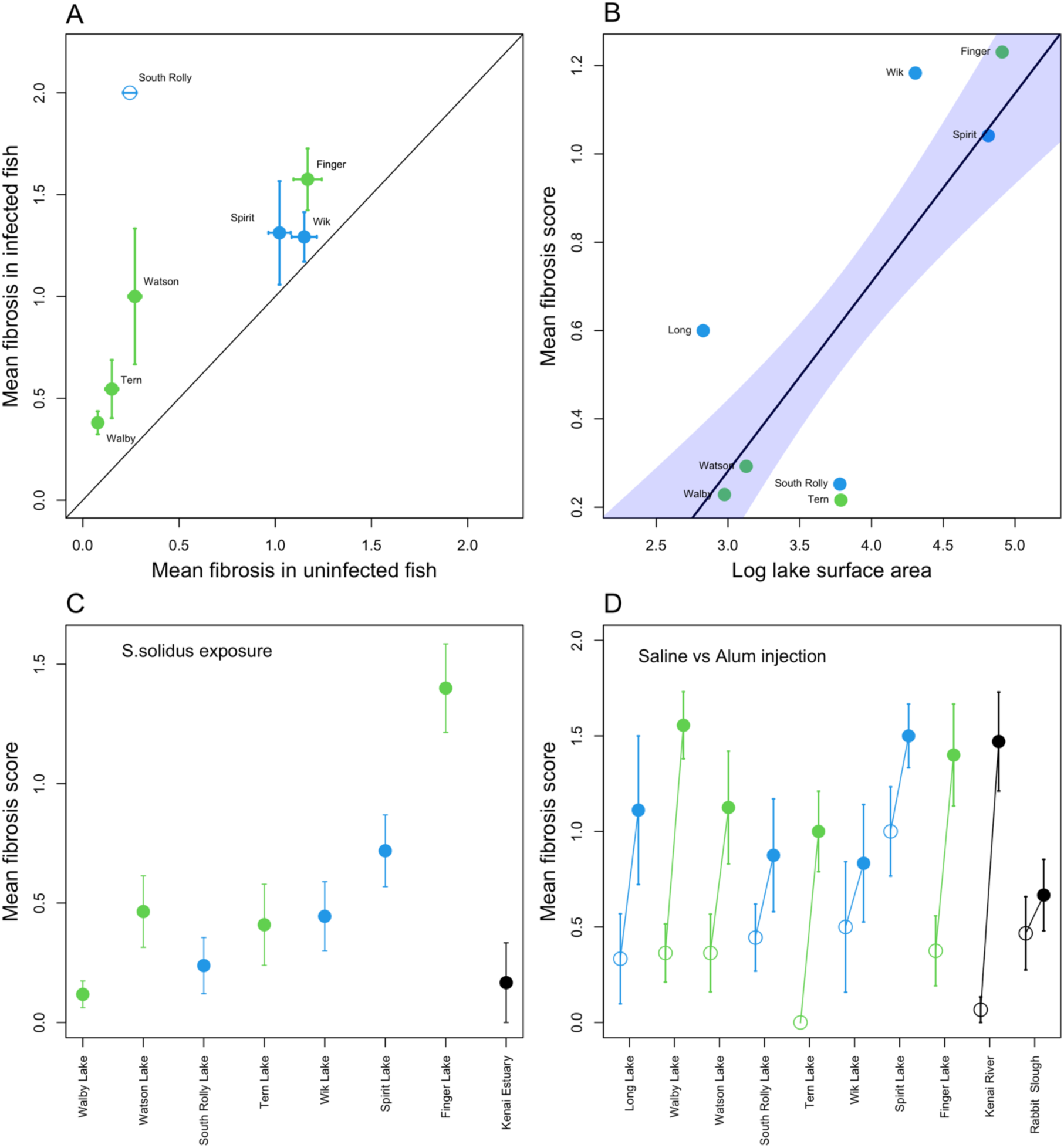
Peritoneal fibrosis in stickleback is induced by *S. solidus* infection, but this response varies heritably among source populations. (**A**) Within each source lake, infected individuals on average have more severe fibrosis than uninfected individuals (linear model infection effect F_1,1791_=11.1, P=0.0008, lake effect F_8,1791_=72.3, P<0.0001). The magnitude of this difference varies among lakes (lake*infection interaction F_7,1791_=2.4, P=0.0205). Data points are color coded by ecomorphologically defined benthic (green) and limnetic (blue) populations, with standard error bars. Figure S4 confirms this effect also holds in recipient lakes. **(B)** Fibrosis is more severe in larger source lakes where stickleback tend to eat more copepods (r=0.826, t=3.59, P=0.012). **(C)** Lab-raised stickleback initiate fibrosis when experimentally fed *S. solidus*-infected copepods (means with standard errors shown), but not when fed uninfected copepods. The magnitude of this fibrosis response varied among source populations (F_7,196_=10.3, P<0.0001), as well as by tapeworm strain (fig. S2). Lakes are ordered from small (left) to large (right). The fibrosis response to cestode exposure increases with lake area (r=0.851, P=0.032). A low-fibrosis marine population (Kenai Estuary, black point) is included to represent an ancestral character state. **(D)** Lab-raised fish from the source populations also initiate fibrosis in response to alum (filled circles) 35 days post injection relative to saline-injected controls (open circles) (injection treatment F_1,186_=8.9, P=0.0032), though this response does not differ significantly among freshwater populations (Population effect F_7,128_=1.3, P=0.24; Population*treatment interaction F_7,128_=0.83, P=0.57).

In theory, host-parasite interactions may lead to stable eco-evolutionary equilibria. However, newly founded populations would be displaced from such equilibrium and should show fluctuations in parasite prevalence as host immunity evolves (*8*). Monitoring infection rates for five years confirmed these expectations: infection rates were persistently different among source populations (Fig. 3A), but destabilized in founded populations (Fig. 3B). In source populations, infection rates differed among lakes (81.6% of binomial GLM explained deviance), with little temporal variation (year and year*lake interaction effects explained 9.7% and 8.7% of deviance; all P<0.0001). In recipient lakes, infection rates fluctuated strongly between years (lake*year interaction explained 66.9% of variance, population 13.9%, year 19.1%, all P < 0.002). Infection rates exhibited negative temporal auto-correlations: lakes with high infection rates one year were rarely infected the next (Fig. 3B). In 2019, benthic founders began with a higher starting infection rate, but the next year infection rates were higher in lakes with limnetic pool fish (t=2.77, P=0.0323). By 2023 infections were again higher in lakes receiving benthic founders (t=-2.011, P=0.0901). Newly founded populations thus experienced destabilized infection rates. In addition, our experiment confirms the ‘enemy release’ hypothesis, which posits that newly founded populations experience reduced parasitism (*3*, *27*); on average the recipient lakes exhibit 77% lower infection rates compared to native source lakes (fig. S3).

**Figure 3.**
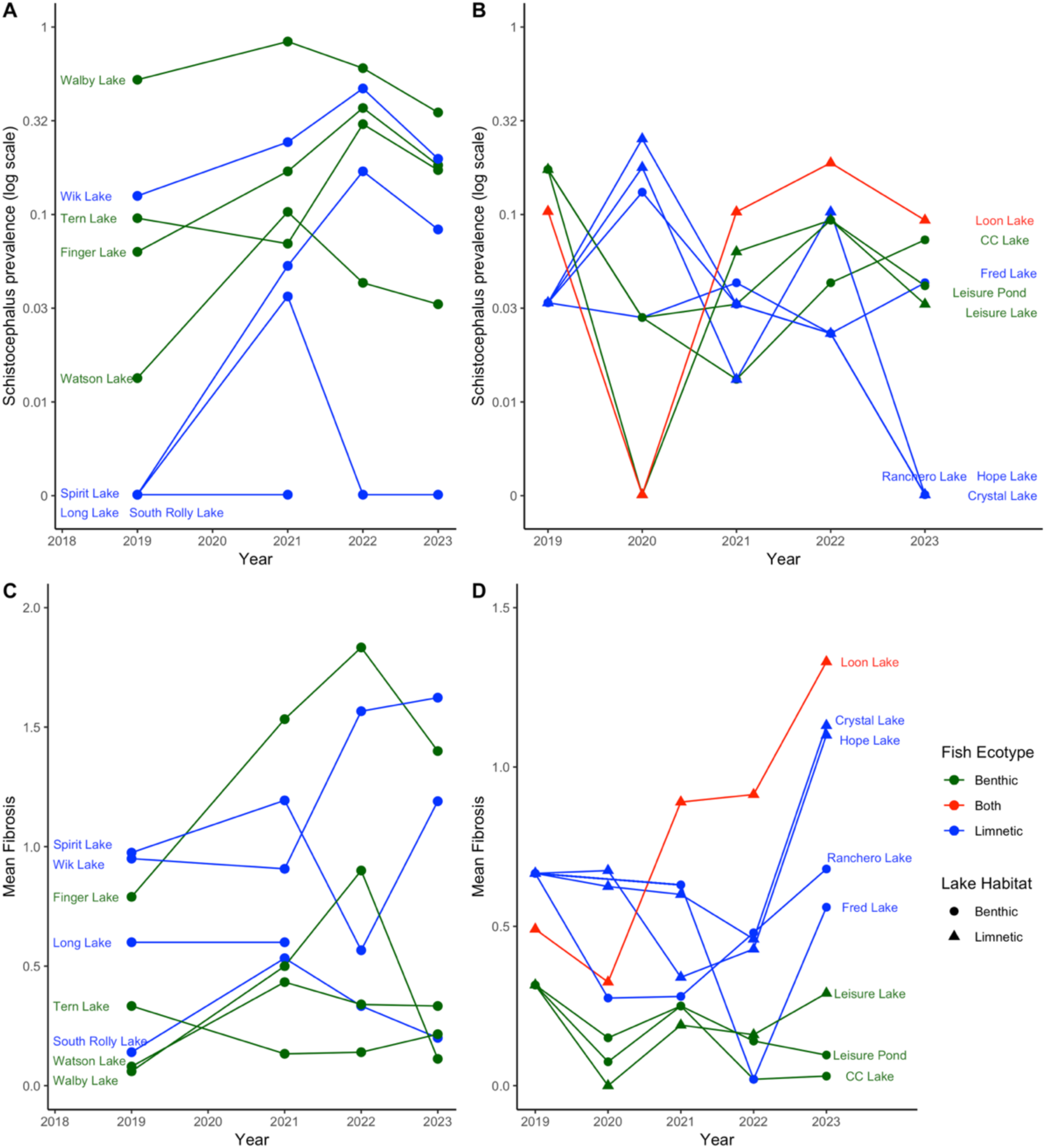
Temporal dynamics of *S. solidus* infection prevalence and fibrosis severity over five years in source and recipient lakes. Points are color coded by source lake fish ecotypes native to, or introduced into, a given lake (green for benthic and blue for limnetic; red for Loon Lake which received both founder pools). Symbols distinguish larger recipient lakes (triangles) and smaller recipient lakes (circles). **(A)** Infection rates varied stably among source lakes (binomial GLM, lake Deviance=378.4, df=7, P<0.0001), with a relatively weak effect of time (Year Deviance=45.0, df=3, P<0.0001; Lake*Year Deviance=50.4, df=19, P=0.0001). **(B)** Infection rates in experimentally founded lakes varied unstably through time (lake Deviance=38.9, df=8, P<0.0001; Year Deviance=33.6, df=3, P<0.0001; Lake*Year Deviance=133.9, df=22, P<0.0001). **(C)** Fibrosis varied substantially among source lakes (F_8,1780_=75.0, P<0.0001, 78% explained variation), with comparatively weak but still highly significant effects of year (F_3,1780_=14.8, P<0.0001, 6%), and a lake*year interaction (F_19,1780_=6.66, P<0.0001, 16%). **(D)** Fibrosis severity diverged among lakes (F_8,2480_=43.2, P<0.0001, 64% explained variation), with significant effects of year (F_3,2480_=26.7, P<0.0001, 15%), and a lake*year interaction (F_20,2480_=5.5, P<0.0001, 21%).

Source populations exhibit stable differences in the severity of peritoneal fibrosis (Fig. 3C), with lake contributing 78% of explained variance (15% for year, 7% for lake*year interaction). In contrast, fibrosis diverged among the newly-founded populations (Fig. 3D), leading to increasing between-lake variance (64% of variance attributed to lake, 15% for year, 21% for lake*year, all P<0.0001). As predicted, this among-lake variation reflects heritable effects of founder type, and effects of local lake habitat. Populations descended from limnetic source lakes inherited a greater propensity for fibrosis (3.1-fold more severe than benthic-founded populations (Fig. 3D; F_2,33_=21.2, P<0.0001). This difference increased over generations (year*ecotype-pool interaction P=0.007), consistent with the expectation that populations predisposed to consume limnetic prey would evolve stronger fibrosis. Fibrosis was also 1.8-fold higher in larger recipient lakes (Fig. 3D, F_1,33_=14.4, P=0.0006, controlling for input ecotypes), consistent with the *a priori* expectation that fish in larger lakes encounter more *S.solidus*, inducing fibrosis more. The fibrosis difference between small benthic and large limnetic recipient lakes grew larger over generations (year*lake interaction P=0.009).

Eco-evolutionary theory suggests that changes in infection rates should be coupled with immune trait evolution. High infection rates should select for stronger immune responses, which feedback to reduce infections (*19*, *28*, *29*). When infections are rare, selection favors loss of costly defenses, renewing opportunities for the parasite. Our time series data confirms there were coupled changes in immunity and infection. In recipient lakes, higher infection prevalence is correlated with stronger fibrosis (P=0.039). This correlation was strong initially (r=0.85 in 2020) but weakened over successive years (Fig. 4A, 2021: r=0.63; 2022: r=0.60; 2023: r=-0.11; year*prevalence interaction, P=0.035). By 2023, several of the most fibrotic populations had few surviving tapeworms (fibrosis is irreversible, persisting after failed infection attempts). This progressive decoupling of infection and fibrosis can occur if high-exposure populations evolved a strong fibrosis response that subsequently reduced infection rates. Confirming this explanation, recipient lakes with high fibrosis in one year, tended to exhibit a stronger decline in *S.solidus* infection rates (Fig. 4B), though this effect varied between years (P<0.0001). For instance, recipient lakes with higher mean fibrosis in 2020 experienced a stronger drop in infection prevalence from 2020-2021(r=-0.77, P=0.042). The same trend held in 2022 (r=-0.65, P=0.079), though not 2021 (r=0.05, P=0.901). These results confirm the prerequisite for eco-evolutionary dynamics: infection promotes fibrosis (fig. S4), which then limits infection. This negative feedback is confirmed by a negative temporal auto-correlation in infection rates: in a given year, recipient lakes with high tapeworm prevalence exhibited a subsequent decline in infection rates the next year (Fig. 4C, prevalence change depended on prior prevalence F_1,24_=84.1, P<0.0001; year F_3,24_=1.8, P=0.181, and prior prevalence * year interaction F_3,24_=5.6, P=0.004). In contrast, there is no significant temporal auto-correlation within the source lakes from 2019-2023 (t=-0.63, P=0.538, fig. S5).

**Figure 4.**
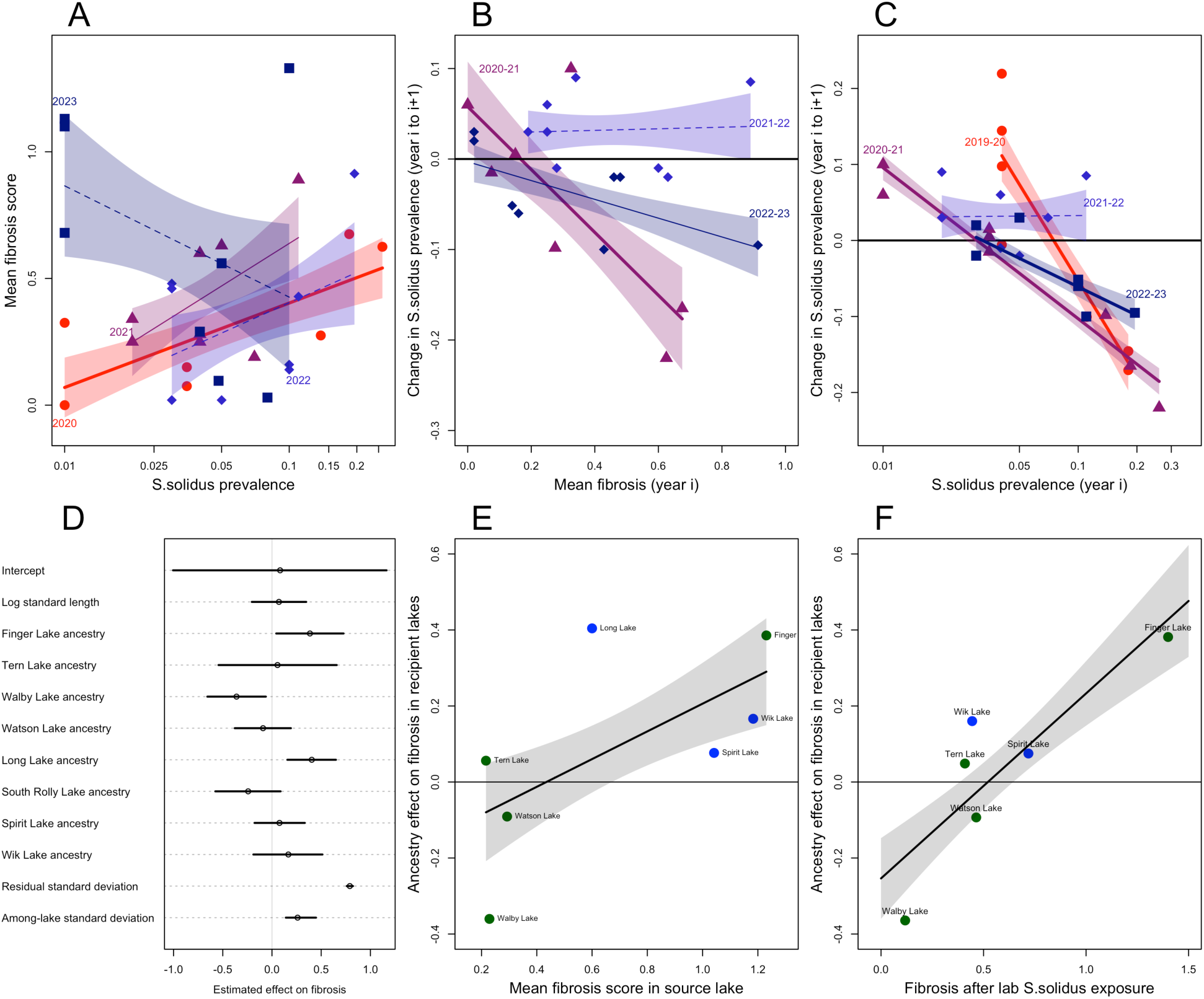
In the recipient lakes, temporal changes in *S. solidus* prevalence are linked to heritable changes in their hosts’ fibrosis response. (**A**) Overall, fibrosis tended to be higher in recipient lakes with higher infection prevalence (2023 being the non-significant exception). In 2020 fibrosis was strongest in lakes with the highest infection prevalence (r=0.85, P=0.015), but this correlation weakened each subsequent year (2021 r=0.63, P=0.093; 2022 r=0.60, P=0.116; 2023 r=-0.109, P=0.779). Each regression line with shaded confidence interval (and point shape) is for a separate year. Thick solid lines are statistically significant (P < 0.05), thin solid lines marginally so (P < 0.1), and dashed lines non-significant. **(B)** Lakes with more fibrosis in a given year exhibited a larger decrease in tapeworm prevalence the following year (note, 2019 fibrosis was not measured so the 2019-2020 contrast is omitted). Each between-year comparison is represented by a different-colored trendline with shaded confidence interval and different color/shape points. A horizontal line is provided to indicate zero change. **(C)** Tapeworm prevalence exhibits negative temporal auto-correlation between most years. With the exception of 2021-2022, lakes with high prevalence in one year had reduced prevalence in the next (and vice versa). Each regression line (with 1 s.e. confidence intervals) is a between-year comparison. See Fig. S5 for an equivalent within the source lakes. **(D)** Fibrosis exhibits heritable variation within the admixed recipient lake populations: individuals fibrosis score depends on their source lake ancestry. We plot Bayesian posterior effect means and 93% credibility intervals. Fish with greater proportional ancestry from Finger or Long Lake had stronger fibrosis, and Walby and South Rolly Lake ancestry reduced fibrosis. Consistent with a priori expectations, these ancestry effect estimates from the experimental recipient lakes are positively correlated with **(E)** the severity of fibrosis in the original source lakes, and **(F)** the strength of fibrosis in lab-raised fish following cestode exposure (r=0.854, P=0.0303).

Despite the rapid timescale of coupled changes in recipient lake infection and fibrosis, several lines of evidence suggest that these involve evolution of heritable immune traits, generating eco-evolutionary dynamics. Our experimental design lets us partition genetic versus environmental effects on traits because we replicated two genetically diverse founder pools across both small and large lakes. We used a SNP array on 2022 recipient lake samples, to determine each F2 individual’s proportional ancestry. Controlling for recipient lake and fish size, fibrosis is higher in individuals with more Finger or Long Lake ancestry and reduced by Walby Lake ancestry (Fig. 4D). Ancestry also affected *S. solidus* infection rates (fig. S6). Fibrosis was yet again a plastic response to infection, higher in infected individuals (P=0.0001, fig. S4). However, there was heritable variation in the magnitude of this plastic response (infection*ancestry: P=0.0034). Fish with more ancestry from low-fibrosis Walby Lake were less responsive to infection, whereas high-fibrosis Finger Lake ancestry conferred stronger response (fig. S7). These estimates of fibrosis heritability are confirmed by two separate datasets. First, source lakes with higher fibrosis have recipient-lake descendents with higher fibrosis (Fig. 4E, r=0.626, P=0.066). Second, ancestry effects on fibrosis were correlated with lab-raised sticklebacks’ responses to *S.solidus* exposure (Fig. 4F, r=0.854, P=0.0303). This consilience of several lines of evidence confirms there is heritable variation in fibrosis among source lakes and within recipient lakes. Therefore, the divergence in fibrosis between recipient lake populations (fibrosis increasing in limnetic-pool and limnetic-habitat lakes) likely represents eco-evolutionary dynamics.

Our replicated whole-lake experiment provides allowed us to observe the earliest stages of host-parasite eco-evolutionary dynamics in newly founded populations. We are able to directly describe the joint dynamics of infection and a key immune trait, peritoneal fibrosis, in the first generations after population founding (fig. S8). Experimentally confirming the ‘enemy release hypothesis’ (*3*), *S.solidus* prevalence was reduced in newly founded lake populations, compared to their source lakes. Following this initial release, we observed large swings in infection rates in the first generations after new populations are founded: heavily infected populations exhibited stronger fibrosis, which then reduced infection rates, leading to negative temporal auto-correlation in parasite prevalence. Over several years, these fluctuations decayed as reintroduced populations diverged in their fibrosis phenotype: stronger fibrosis emerging in populations that inherited limnetic genotypes, or inhabited limnetic lakes, where *S.solidus* exposure is more likely. These results provides a unique demonstration that large changes in parasitism and immunity can occur in just a few generations and can lead to rapid among-population divergence in ecology and immunity. These rapid changes would be overlooked by observational studies of already-established invasive species. Because parasitism has a large impact on population viability, the eco-evolutionary dynamics documented here may play a key role in the early success or failure of invasive species (*13*, *30*), geographic range expansion under climate change, and species reintroductions for conservation.

## Acknowledgments

This research was made possible with support from the Alaska Department of Fish and Game, with particular thanks to Robert Massengill. Field work benefitted from assistance from a large group of field workers (full list in Sup. Mat).

## Author contributions

Conceptualization: APH, DIB, RDHB, KMM, CLP, JNW, NCS

Methodology: DIB, RDHB, APH, KMM, CLP, JNW, NCS

Investigation: DIB, RDHB, LE, APH, EVK, ÅJL, KMM, CLP, KS, KTS, ART, CW, JNW, NCS

Visualization: DIB, EC

Funding acquisition: DIB, KMM, CLP, JNW, NCS

Project administration: APH, DIB

Supervision: DIB, JNW, NCS, KMM

Writing – original draft: DIB

Writing – review & editing: All authors

## Data and materials availability

All data files and R code required to reproduce the results in this paper have been archived at Figshare.org 10.6084/m9.figshare.26093566.

## Supplementary Materials

Materials and Methods

Supplementary Text

Figs. S1 to S8

Tables S1 to S5

## The Supplementary Materials

### Materials and Methods

#### Experimental design

In 2018, the Alaska Department of Fish and Game used rotenone to eliminate an invasive species of fish (northern pike) from nine lakes on the Kenai Peninsula, thereby eliminating all fish. Surveys in spring 2019 confirmed the lakes remained fishless. In May-June 2019, we used minnow traps to collect stickleback from eight lakes with extant stickleback populations (“source lakes”). We selected populations that differed in body shape, spanning a well-known ecomorphological gradient from benthic to limnetic ecotypes (*29*, *32*, *40*, *41*).

Four lakes contain relatively limnetic stickleback populations, characterized by more fusiform body shape typically associated with feeding on zooplankton in midwater (including copepods that transmit *S.solidus* tapeworms). Four source lakes contain relatively benthic stickleback populations, with deeper bodies and larger jaws associated with eating shallow-water benthic invertebrates. Note, however, that these ecotypes form a continuum, rather than representing two discrete morphs (*40*), and even within lakes individuals vary along this benthic-limnetic axis (*41*). In general, stickleback in larger lakes tend to be more limnetic (and benthic in small lakes) (*29*). However, in the eight source lakes used for this experiment, lake size and ecotype are not significantly correlated (e.g., the largest lake, Finger Lake, is morphologically relatively benthic).

We pooled approximately equal numbers of fish from the four benthic source lakes to create a benthic pool, and likewise created a limnetic pool from the four limnetic lakes. The benthic pool was distributed into four fishless “recipient lakes” (two larger and two smaller lakes). The limnetic pool was distributed into four other recipient lakes (two larger and two smaller). The result is a factorial design with two replicate lakes for each combination of benthic (or limnetic) source pool, and benthic (or limnetic) recipient lake. A ninth recipient lake received both the benthic and limnetic pools, mixed together. The numbers of transplanted fish are listed in table S1, and the experimental design is illustrated in Fig. 1A. One of the recipient lakes, G Lake, failed to establish a population and is not considered further here. The transplants were conducted with IACUC approval (McGill University AUP 2000-4570) and permits from the State of Alaska (Aquatic resource permits SF2019-085, P-19-005) and Kenai National Wildlife Refuge (2019-Res-AHendry-6576). For more details on the experimental design, see citation (*1*). A complete acknowledgement list of field assistants is provided at the end of this supplementary Methods document.

#### Sampling

Each year we sampled stickleback to determine parasite prevalence and fibrosis severity. Minnow traps were placed overnight along the shoreline of each lake. All fish captured from each lake were pooled and 100 randomly sampled individuals were euthanized in MS-222 and retained for data acquisition. In 2019 we sampled stickleback from the 8 source lakes just prior to introductions began. Infection and fibrosis prevalence for the new transplanted populations in 2019 are inferred as a weighted average of the prevalences of the source lakes used to colonize each recipient lake. We note that sampling error and biased mortality might alter the actual infection rate of founders. In 2020 (due to pandemic limitations) we sampled only the recipient lakes. In 2021, 2022, and 2023 we sampled both source and recipient lakes. A few source lakes yielded low capture rates in later years, possibly due to increasing abundance of invasive pike in those locations. Sample sizes are summarized in table S2. Lethal sample collection from source and recipient lakes was conducted with IACUC approval (University of Connecticut A22-006) and permits from the State of Alaska (SF2020-103d, P-21-012, SF2022-043d, SF2023-030d).

#### Phenotypic measurements

Fish were photographed, weighed, and standard length measured with calipers. They were then dissected to count *S. solidus* parasites (a strict specialist on threespine stickleback). Sex was determined by visual inspection of gonads. Fibrosis was scored on an ordinal scale from 0 to 4 following protocols in (*34*). A score of zero means no fibrosis, the organs move freely separate from each other and from the body wall. A score of 1 denotes moderate thread-like connections between organs, typically the liver to intestines. A score of 2 indicates extensive connections between organs that can be separated by force. A fibrosis of 3 is when the organs are fully encased in a cocoon of fibrosis and cannot readily be separated without damage to the organs, and the organs are attached to the body cavity by fibrotic threads. A score of 4 indicates the body wall cannot be separated from the organs without tearing the muscle tissue or organs. The visual fibrosis scoring is highly repeatable: independent observers’ scores are highly correlated (r>0.95).

#### Ancestry Inference

To infer ancestry of fish from the recipient lakes, we genotyped individuals for SNPs diagnostic of each source population. Using PoolSeq data from the source populations, derived from (17), we selected 24 SNPs per source population that were unique to that population (table S3). We selected SNPs to maximize the frequency of the unique alleles in the respective source populations, while filtering out SNPs with low read numbers (more than 1.5 SDs fewer than the mean read number) and avoiding effects of linkage by ensuring that every chromosome (excluding the sex chromosome) had at least one SNP. We designed two Fluidigm SNPtype assays, with SNPs from the benthic and limnetic source populations respectively. To validate the efficacy of these assays in correctly inferring ancestry, we genotyped 16 individuals from each source population from samples collected by (20) in 2018, all of which were assigned to the correct population based on the genotyping results. This trial run also identified SNPs that were unsuccessful or that underperformed (were present in the trial fish in much lower frequency than expected from the PoolSeq data), which were omitted from subsequent analyses, leaving 20 SNPs per population on average. The final list of SNPs, 158 in total with an average allele frequency of 81%, can be found in Table S3.

Using these assays, we genotyped 95-97 fish from each of the recipient lakes in 2021, extracting DNA using a phenol-chloroform extraction protocol on tissue (caudal fin) samples that were taken shortly after the fish were euthanized. To infer the ancestry of fish from the genotyping data, we computed an ancestry “score” for each source population for each individual. This score is based on the number of unique SNPs identified from each source in that individual, weighted to account for differences in the number of SNPs included for each source and the average frequency of those alleles in the source populations. Finally, the proportional scores for each source population within each individual fish were rounded to the nearest fraction deemed possible according to the maximum potential generations, while ensuring the proportions always summed to 1. We assumed the fastest generation time to be one year, making the maximum generation in 2021 the F2, so the proportions were rounded to the nearest multiple of 0.25. To confirm the accuracy of these inferences with this number of SNPs, we simulated analogous scenarios in R. In the simulated F1 generation, we have an error rate of 0.005% in 100 simulations, and in the F2, we have an error rate accuracy of 6.5%. For the individuals whose ancestry we mis-infer, we still identify the correct set of ancestral populations (but infer the wrong proportions) about half the time, leaving just 3.5% of F2 individuals where we fail to identify one of ancestral populations.

#### Breeding of fish for lab-based experiments

In 2021, source lake stickleback, and control marine ancestors (Rabbit Slough, Kenai Estuary), were bred following previously described *in vitro* fertilization protocols (ref) (Alaska Aquatic Resource Permit P-21-008). In brief, we collected gravid fish via unbaited minnow traps placed overnight along the shoreline. After euthanizing gravid fish in MS-222, eggs were stripped from females and combined with macerated testes in a petri dish. Fertilized eggs transported to the University of Massachusetts Lowell and the University of Wisconsin-Madison for rearing.

#### Injection assays of fibrosis variation between source populations

Fertilized eggs were reared to maturity at the University of Massachusetts. Following hatching, stickleback were grouped by family and housed at 17℃, with a 18:6 hr light:dark cycle in a modified zebrafish recirculating system. Two-year-old fish were injected intraperitoneally with either PBS (0.9x endotoxin free PBS), alum (1% AlumVax Phosphate, OZ Bioscience AP0050). Two families from each population were injected, with the exception of Finger Lake which had smaller family size so three families from this populations were used. Within each family, fish were divided evenly among treatments (Table S4). Prior to injection fish were anesthetized in MS-222 (50 mg/mL, pH 7.4) until non-reactive to stimuli. Fish were placed on a paper towel covered sponge, soaked in system water and the fish’s eyes and opercular flaps were covered with a wet paper towel. The peritoneal injections were administered into the peritoneal cavity on the left side. Fish were immediately returned to a recovery tank and monitored until alert. 35 days post injection fish were euthanized in MS-222 (500mg/ml, pH 7.4) for at least 5 minutes and fibrosis was scored using the previously described scoring system. Animal husbandry and injection experiments were conducted under approved Institutional Animal Care and Use Committee protocol 21-10-07-Ste.

#### Infection assays of fibrosis variation between source populations

*S. solidus* were collected from infected threespine stickleback from Walby, Finger, and Tern Lakes in Alaska, as well as Lake Kjerringøy in Norway. Fertilized eggs were harvested from the cestodes using the methods in (*20*). The cestode eggs were incubated in the dark at 18°C for 3-7 days before being exposed to light to induce hatching. *Acanthocyclops robustus*, a cyclopoid copepod, were fasted for 24 hours prior to being exposed to hatched cestode coracidia. A subsample of copepods from each batch exposure was dissected after 14 days to determine the average infection rate. To expose stickleback to the infected copepods, food was withheld from the fish for 24 hours to encourage copepod ingestion and each fish was placed in a small Tupperware container with a tube to oxygenate the water. The estimated infection rate of *A. robustus* was used to determine per fish exposure rates.

Lab data for fibrosis represent a pool of two related experiments. In the first experiment (labeled experiment “A” in Table S5 and in supplemental data), individual stickleback were placed in a Tupperware with 500mL of tank water and 10-20 infected copepods so that every fish was exposed to a minimum 10 *S. solidus*. The fish were returned to their home tank after remaining in the container overnight and the water was filtered through a 250μm sieve to confirm that the copepods were consumed. Approximately 41-48 days after exposure fish were euthanized in MS-222 (500mg/ml, pH 7.4) for at least 5 minutes followed by pithing. The fish were then dissected and fibrosis was scored using the previously described scoring system.. We also fed uninfected copepods to stickleback to rule out an effect of copepod consumption on fibrosis. In this case, ∼2000 uninfected copepods were introduced to tanks of 15-20 stickleback that had been starved for 24-hours, and fish were dissected 13-15 days later. Sample sizes are presented in table S5.

In the second experiment (‘B’ in table S5), fish were exposed to approximately 7-8 Walby Lake cestodes with a few small modifications. First, fish were placed in population-specific, mixed family tanks at least 28 days before exposure (4 Spirit families, 3 Wik families, 3 Watson families, and 3 Finger families). They were then acclimated to 18°C and, as part of the separate experiment, additional data was collected from each fish. Specifically, they were subjected to more handling than in experiment A and fasted for 48 hours prior to exposure to infected *A. robustus*. We dissected fish and scored fibrosis 30 days post cestode exposure. The preceding methods were conducted with University of Wisconsin-Madison IACUC approval (protocol number L006460-A04).

#### Analysis

All of the following analyses were conducted using the R statistical language, version 2023.06.1+524 (*42*). R code is available on the data and code repository accompanying this paper.

##### Are the source populations genetically divergent?

We re-analyzed previously generated genomic allele frequency data derived from PoolSeq libraries from each of the eight source populations (*21*). For outgroups, we also used PoolSeq data from two anadromous-marine populations (Rabbit Slough in Alaska, and Sayward Estuary in British Columbia). From these allele frequencies we calculated pairwise F_ST_ between populations, first for each SNP, then averaged F_ST_ across SNPs to generate an overall measure of allele frequency divergence between each pairwise comparison of populations. We used the R package *ape* v5.7-1 (*43*) to generate a neighbor joining tree from the F_ST_ distance matrix.

##### Do source populations differ in infection prevalence and fibrosis severity?

We began by calculating the prevalence and mean intensity of *S. solidus* infection in each lake, in each year. Prevalence is the proportion of fish with infections present (regardless of the size or number of tapeworms), with confidence intervals calculated using the R package *exactci* v1.4-4 (*44*). Intensity is the mean number of tapeworms per individual fish. Focusing on source lake data from 2019 when the experiment was initiated, we used a binomial general linear model (GLM) to test whether prevalence differed among source lakes. We used a Poisson GLM to test for intensity differences among lakes. In both analyses we used fish size (standard length) and sex as covariates. The differences among source lakes in 2019 will reflect the different inputs of tapeworms into the recipient lakes. We then repeated the analysis with all years for which we have source lake data (excepting 2020, when the COVID pandemic limited our sample collection). We fit a general linear model testing for effect of lake, and year, on either infection prevalence (binomial GLM) or intensity (Poisson GLM). Sex and log length were again used as covariates. Using the estimated population mean prevalences, we used a t-test to determine whether infection rates differed between the four benthic versus limnetic lakes (defined morphologically, (*32*)). We used a correlation test to evaluate whether infection rate is positively correlated with lake size (hectares).

Fibrosis is scored on an ordinal range from 0 to 4, but for simplicity we use linear models to test whether fibrosis intensity depends on source lake, with Type II Sums of Squares in an ANOVA analysis, with permutations used to calculate P values because the dependent variable is ordinal. We first did this for 2019 alone (to reflect the likely phenotypes of transplanted fish), and pooling all years for a given lake. Then we did a linear model testing for effect of lake, year, and lake*year interactions (with fish length and sex as covariates). Next, treating lakes as the level of replication, we used a t-test to compare mean fibrosis severity between benthic and limnetic populations. We used a correlation test to compare mean fibrosis to lake size, and to prevalence or intensity.

##### Are the source population differences in fibrosis heritable?

We tested for differences in fibrosis among laboratory-raised stickleback from the source lakes. As described above, these fish were bred from wild-caught parents but raised from eggs in the laboratory. They were then either injected (saline controls, alum, or NPCGG in alum; at the University of Massachusetts Lowell), or experimentally exposed to *S.solidus* (at University of Wisconsin-Madison). For the injected fish, we fit a linear model seeking to explain fibrosis severity as a function of treatment contrast (saline vs alum, or saline vs NPCGG), population, and a treatment by population interaction. P values were generated by permutation because the dependent variable residuals are non-normally distributed. A significant population by treatment interaction would denote heritable differences in response to immune challenge. For the cestode-exposed fish, we fit a linear model with fibrosis as a function of fish population, cestode population, and their interaction. We also tested simpler models, omitting the interaction (which was not significant).

##### Does cestode infection induce fibrosis in source lake fish?

Based on laboratory infection experiments and past field samples we expect to see that cestode infection induces fibrosis. To test this observationally, we used a linear model to test whether fibrosis severity (ordinal, 0,1,2,3,4 scores) is correlated with the presence or absence of a tapeworm in each individual fish (or, we repeated this for tapeworm intensity). The linear model included infection, and fish population (a random effect), and an infection*population interaction. Sex and fish length were initially included as covariates, but dropped for lack of explanatory power (based on AIC). Permutations were used to generate P values given the non-normal nature of the fibrosis ordinal score residuals. A positive main effect of infection on fibrosis confirms our expectation. An interaction with source population would indicate that populations differed in their fibrosis response to infection.

##### Do reintroduced populations experience a reduction in infection by S. solidus?

We tested for differences between source versus recipient lakes, separately for each sample year where we have available data (2021, 2022, 2023). Within each year we used a general linear mixed model (glmer in R) with a binomial (for prevalence) or Poisson (for intensity) to test for differences between source and recipient lakes. Lake identity was treated as a random effect. We then tested for an overall effect of source versus recipient lakes using a GLM with both lake type (source/recipient) and year and lake type by year interaction effects.

##### Does infection differ among reintroduced populations differ, and among years?

We used generalized linear models to test whether infection prevalence (binomial) or intensity (Poisson) differ among reintroduction lakes, by year, or as a function of lake by year interactions. Fish sex and length were initially included as covariates. We use planned contrasts within years to test for among-lake variation, and whether that is structured by the lake habitat (e.g., categorical classifications of benthic or limnetic lakes, or lake size as a quantitative metric), or the genotype(s) of fish that were added (benthic pool, limnetic pool, or both). We treat years as a factor rather than numeric variable to avoid any assumption of linear changes over time.

To test for temporal auto-correlations in infection prevalences, we calculated the change in infection rate between each successive year. As a stand-in for initial infection rates in 2019, we calculated the expected prevalence by averaging the relevant source lake prevalences. We then calculated correlations between the change in prevalence between years, versus the previous year prevalence. Equivalent results were obtained by regressing year (i+1) against year i prevalences. We did similar temporal auto-correlation analyses for source lakes, for comparison. Although, lacking 2020 data due to the COVID pandemic we had to use 2019-2021 as one of the time steps.

##### Does fibrosis differ among reintroduced populations differ, and among years?

We used linear regression (with permutation-obtained P values) to test for among-lake and among-year variation in fibrosis in the recipient lakes. We then used estimates of the mean fibrosis for each population in a linear model to evaluate the effects of source lake habitat (benthic or limnetic) and the ecotype of fish introduced (benthic or limnetic).

##### Does ancestry impact infection or fibrosis?

Stickleback sampled from reintroduction lakes in 2021 were genotyped with a SNP array (see above) to determine their proportional ancestry from the eight source lakes. Because each pool of source lakes was introduced to multiple recipient lakes, we can estimate effects of ancestry, and present environment (lake, or lake type) on fibrosis and infection. We used a Bayesian hierarchical linear model to estimate the effect of each source lake (% ancestry) on fibrosis (linear model) or infection (binomial). Lakes were treated as random effects. Fish size (standard length) was included as a covariate. The model was fit with stan in R using the *rethinking* package (*45*), and 93% credible intervals and posterior distribution means were retained. We used correlation tests to compare the mean posterior probability estimates of source lake effects on fibrosis, with other independent measures of the source lakes. In particular, we evaluated whether the effect of source lake ancestry on fibrosis is correlated with infection prevalence or mean fibrosis in the source lakes, lake size, and fibrosis response in laboratory infection trials. We repeated the analyses adding ancestry by infection interaction effects to account for different genotypes responses to infection, and we fit models without ancestry effects. We used WAIC model comparison to select models best supported by the Bayesian analyses.

##### Does cestode infection induce fibrosis in recipient lake fish?

We used a linear model to test whether fibrosis of individual fish (as the level of replication) depends on presence or absence of cestode infection, with lake and year effects, and all two– and three-way interactions. This is an extension of an analysis described above for source lake fish, but replicated in recipient lake fish. We examined the results in greater detail using planned contrasts within each lake, within each year. In the recipient lakes we have the additional benefit of having variance in ancestry (for the 2021 sample). We can therefore test whether ancestry modifies individuals’ fibrosis response to infection. For this, we used linear regression to test whether fibrosis depends on infection status, lake, and ancestry principal component axes (PC1 or PC2), and their interactions. An interaction between infection and ancestry would indicate heritable differences between source lakes in their propensity to respond with fibrosis to tapeworm infection.

#### Extended acknowledgements

This work was made possible by the efforts of a large number of short-term field assistants. The initial 2019 survey of source lake fish was carried out with help from Christopher Peterson, Trey Sasser, Elsa Diffo Tiyao, Kelly Ireland, and Rachel Kramp. The transplants were conducted with help from Hilary Poore, Luis Baertschi, Matt Josephson, Christopher Peterson, Ismail Ameen, Anya Mueller, Chelsea Bishop, Victor Frankel, Allegra Pearce, Matt Walsh, and Michelle Packer. Resampling from source and recipient lakes in later years was done with help from Andrea Roth, Maria Rodgers, Arshad Padhiar, Annika Wohlleben, Kevin Newmann, Kelly Ireland, Emily Kerns, Saraswathy Vaidyanathan, Steve Bezdecny, Rogini Runghen, Sarah Pasqualetti, Abdoleman Nouri, and Benjamin Sulser.

**Fig. S1.**
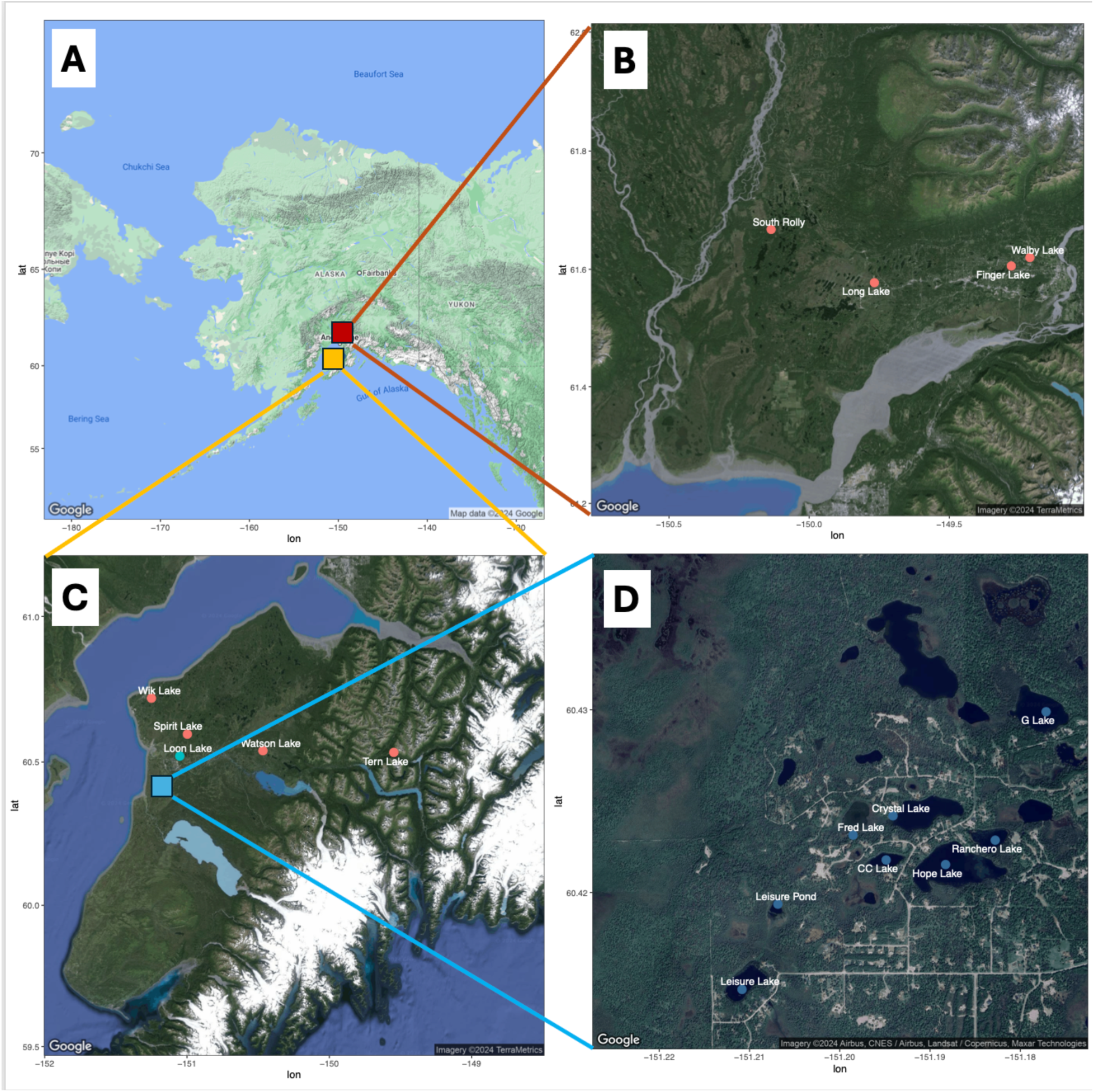
Map of populations used as sources of founders (red points) and fishless recipient lakes (blue points). (**A**) A view of Alaska showing the location of the Mat-Su Valley (red box) and Kenai Peninsula (orange box). **B)** zoom in to four source lakes in the Mat-Su regions. **C)** zoom in to the Kenai Peninsula showing the locations of four source lakes (red points) and recipient lakes (blue). **D)** zoom in to eight of the recipient lakes.

**Fig. S2.**
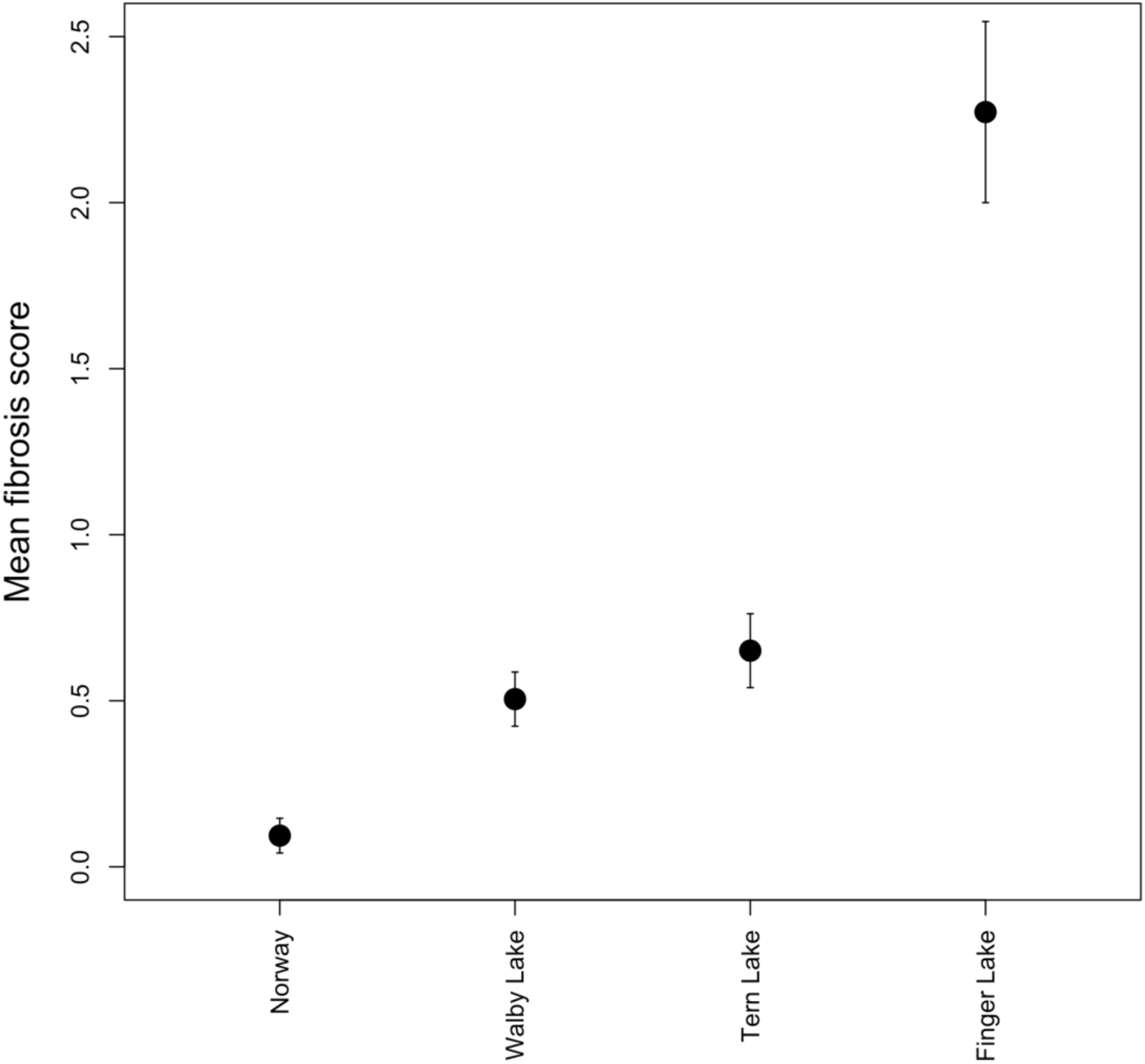
Different tapeworm populations (x axis) induce divergent fibrosis responses in lab-raised stickleback (fibrosis is typically a score of zero in lab-raised fish without an immune challenge). Exposed fish were fed copepods containing procercoids from one of four *S. solidus* source populations. Figure 2B shows that stickleback genotype (source lake) affected the magnitude of fibrosis response. A linear model confirms that fibrosis depends on both stickleback genotype (F_7,196_=10.3, P<0.0001) and tapeworm strain (F_3,196_=8.9, P<0.0001), though there is no detectable fish by parasite genotype interaction effect (F_12,184_=1.2, P=0.2832).

**Fig. S3.**
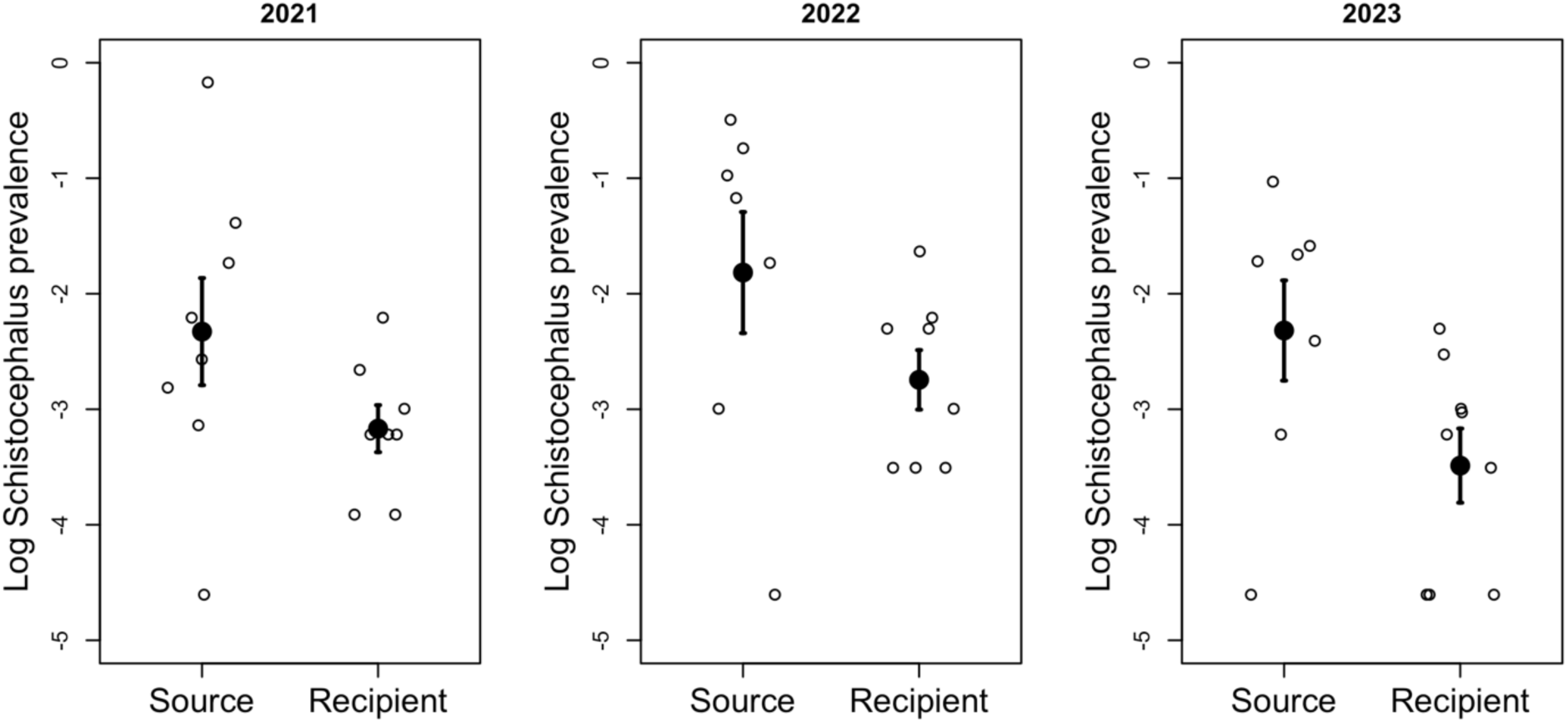
A comparison of *Schistocephalus solidus* prevalence in source versus recipient lakes, in each of the three years in which we have direct comparisons. The reduced infection rate in recipient lakes is consistent with the ‘enemy release hypothesis’. Year effect (treated as a factor rather than linear trend), F_2,41_=1.2, P=0.309; Source/Recipient effect F_1,41_=12.1, P=0.0012; Year*source/recipient interaction effect F_2,41_=0.4, P=0.6801.

**Fig. S4.**
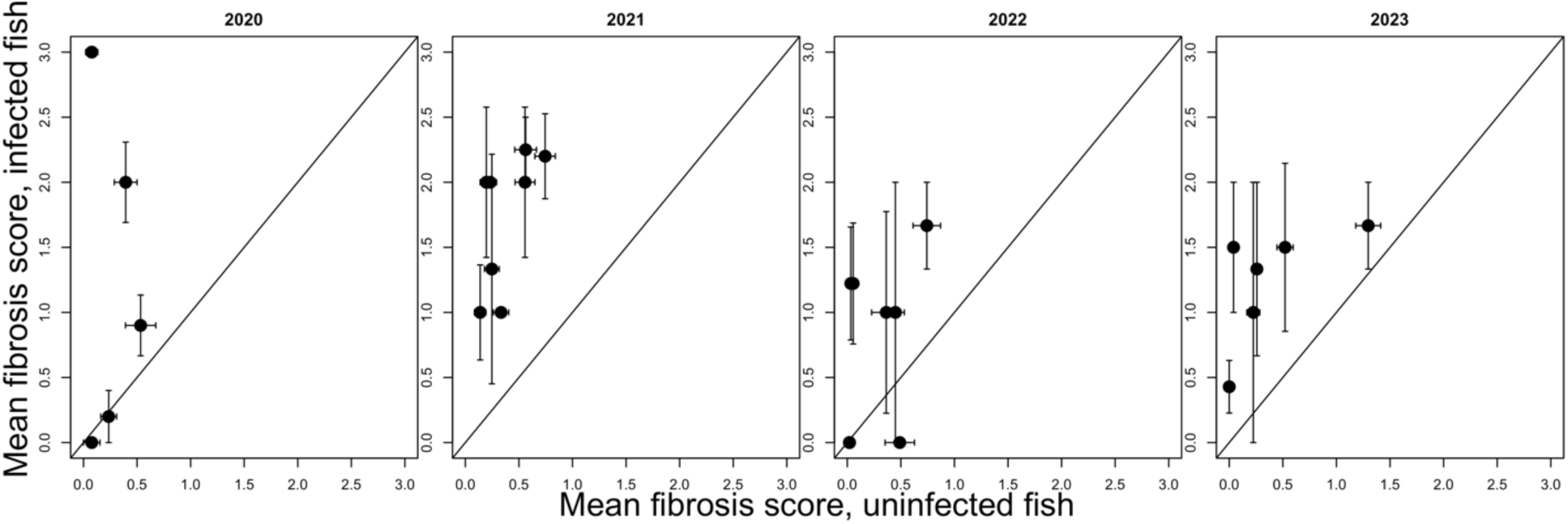
In recipient lakes, individual stickleback with cestode infections have higher fibrosis than uninfected individuals from the same lake. Each panel is a successive year. One standard error confidence intervals are plotted for estimates of mean fibrosis score for each category of fish. Each point represents a single recipient lake. Statistical analysis (Type II ANOVA) revealed a significant increase in fibrosis in response to infection (F_1,2452_=172.11, P<0.0001). This effect recapitulates the source lake result from Fig. 2A. We also observe fibrosis variation among lakes (F_8,2452_=42.6, P<0.0001) and among years (F_8,2452_=34.9, P<0.0001). There is significant variation in this response among lakes (lake*infection interaction F_8,2452_=2.75, P=0.0050), and among years (year*infection interaction F_3,2452_=4.6, P=0.0032). If fibrosis was a purely plastic response to infection that did not evolve over time, we would not necessarily expect changing infection response over time. Therefore, these interaction effects lend support to the inference that fibrosis response to infection is evolving, and diverging between lakes over time (lake*year interaction F_20,2452_=5.58, P<0.0001, lake*year*infection F_15,2452_=1.58, P=0.0712).

**Fig. S5.**
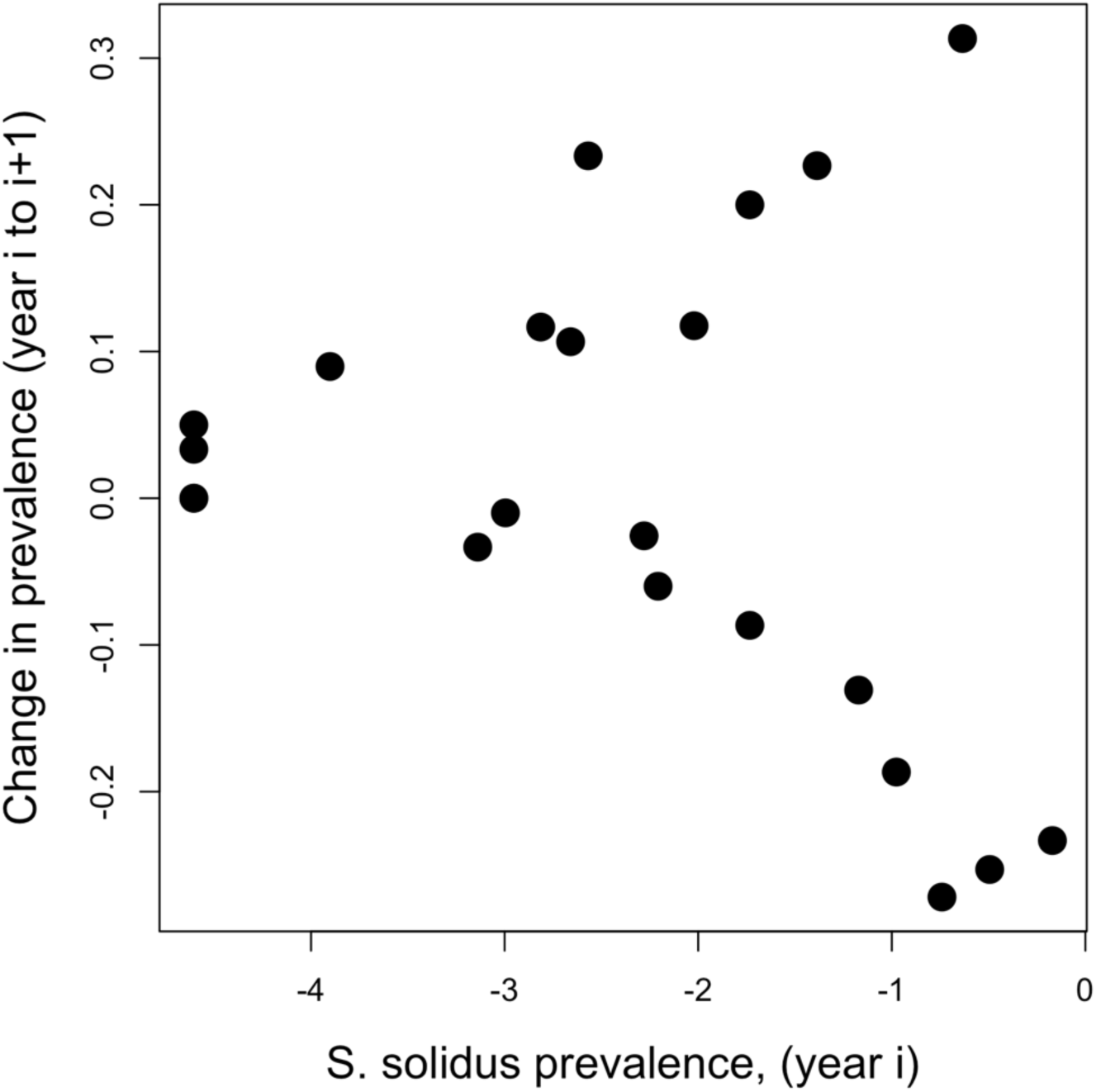
In contrast with Fig. 4C, we found no significant temporal negative auto-correlation within source lakes. Each point is a lake/year combination. There is a significant effect of year (F_2,18_=4.71, P=0.0226) but not prior prevalence (F_1,18_=2.48, P=0.1322).

**Fig. S6.**
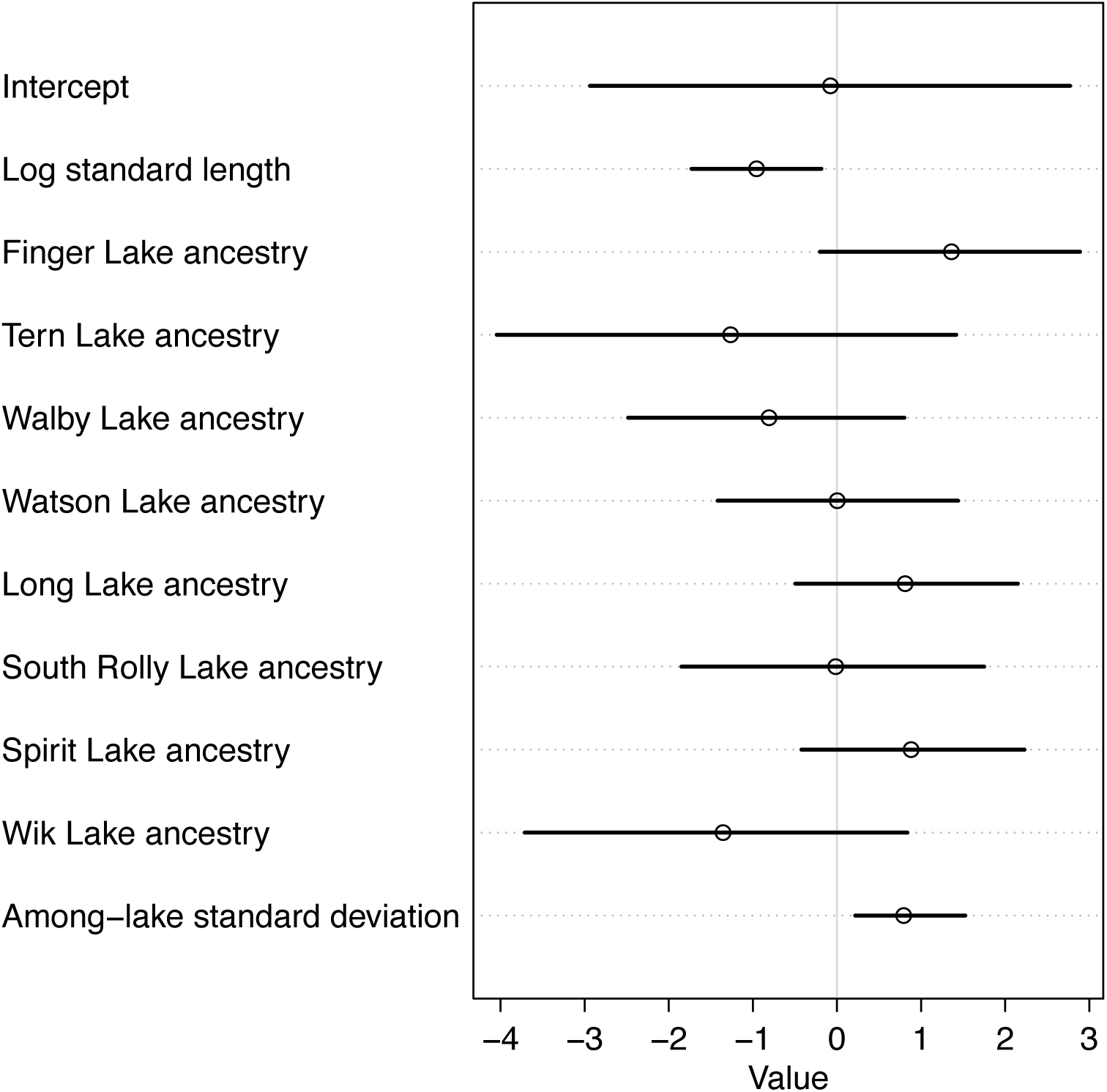
Genetic variation in infection rates within recipient lakes. The figure presents posterior probability means (points) and 93% credible intervals from Bayesian hierarchical linear model analyses of the effect of source lake ancestry, and recipient lake, on *S. solidus* infection probabilities. Effect size estimates indicate which populations contribute increased (positive) or decreased (negative) risk of infection or severe fibrosis. Effect sizes for infection mostly span zero, though there is a clearly positive and non-zero estimate for the among-lake random effect variance.

**Fig. S7.**
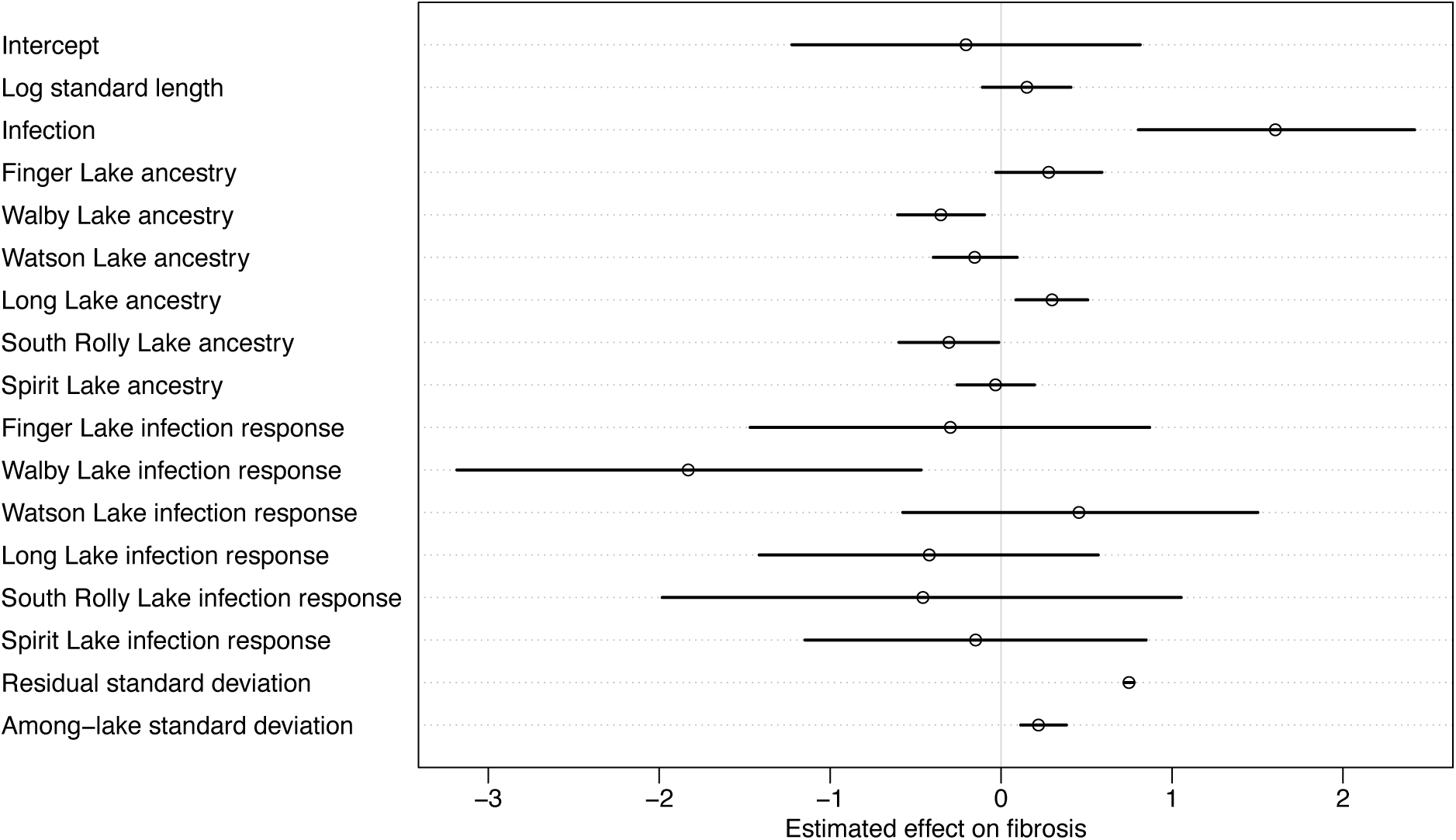
Variation in genotypic responses to infection within recipient lakes. We used a Bayesian linear model to estimate effects of infection, ancestry proportions, and their interactions on within-lake variation in fibrosis, treating recipient lake as a random effect. Here we plot posterior distribution sample means with 93% credibility intervals. Consistent with results reported above (e.g., Fig. S7), there is a positive effect of infection on fibrosis. Baseline fibrosis differs among ancestries. This can reflect genotype-specific responses to prior cestode exposures that failed to generate a detectable infection (e.g., if the fibrosis response successfully eliminated the parasite), because fibrosis persists for many months after a parasite encounter. Thus, the baseline ancestry effects do not entirely represent a parasite-free control. Nevertheless, we detect strong support for additional variation in fibrosis generated by genotype*infection interactions (ancestry infection responses). WAIC comparisons favor models with some of the genotype*infection interactions (85% WAIC weight), over models omitting all genotype-specific responses (15% WAIC weight).

**Fig. S8.**
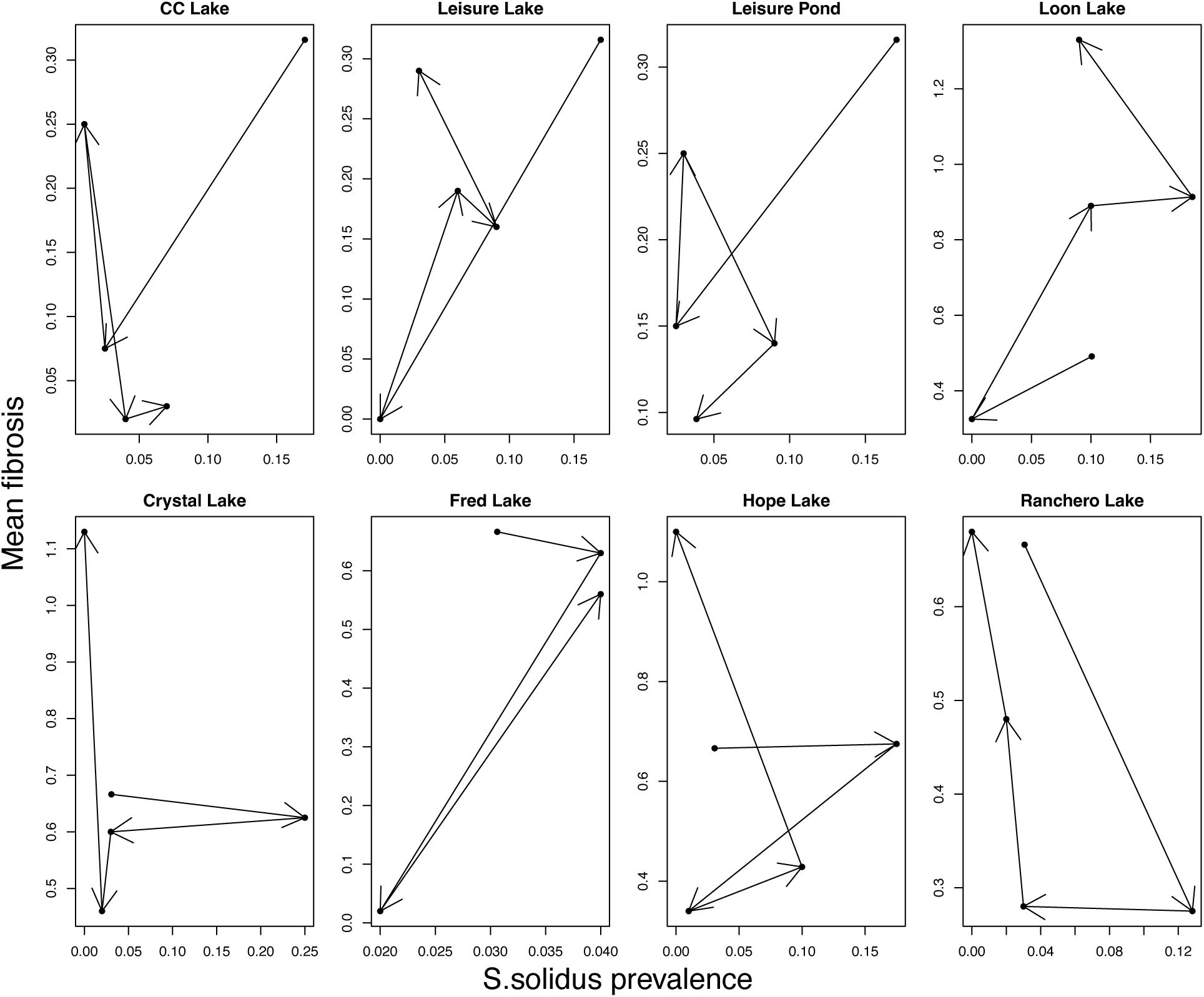
Coupled changes in Schistocephalus solidus infection prevalence, and fibrosis severity, in the eight recipient lakes. Each panel is a single lake population, points represent joint values of infection and fibrosis for a given year. Arrows connect values between years starting in 2019 to 2020, then 2020-21, et cetera. The top row of lakes (except Loon Lake) received benthic pool founders, and experienced an initial decline in infections and fibrosis, followed by an increase in fibrosis. The bottom row of lakes received limnetic pool founders and all experienced initial increases in infection prevalence. Although the exact temporal trajectory of infection and fibrosis differed between populations, this illustration exhibits the tendency for large changes in fibrosis and infection to coincide, consistent with unstable eco-evolutionary dynamics.

**Table S1.**
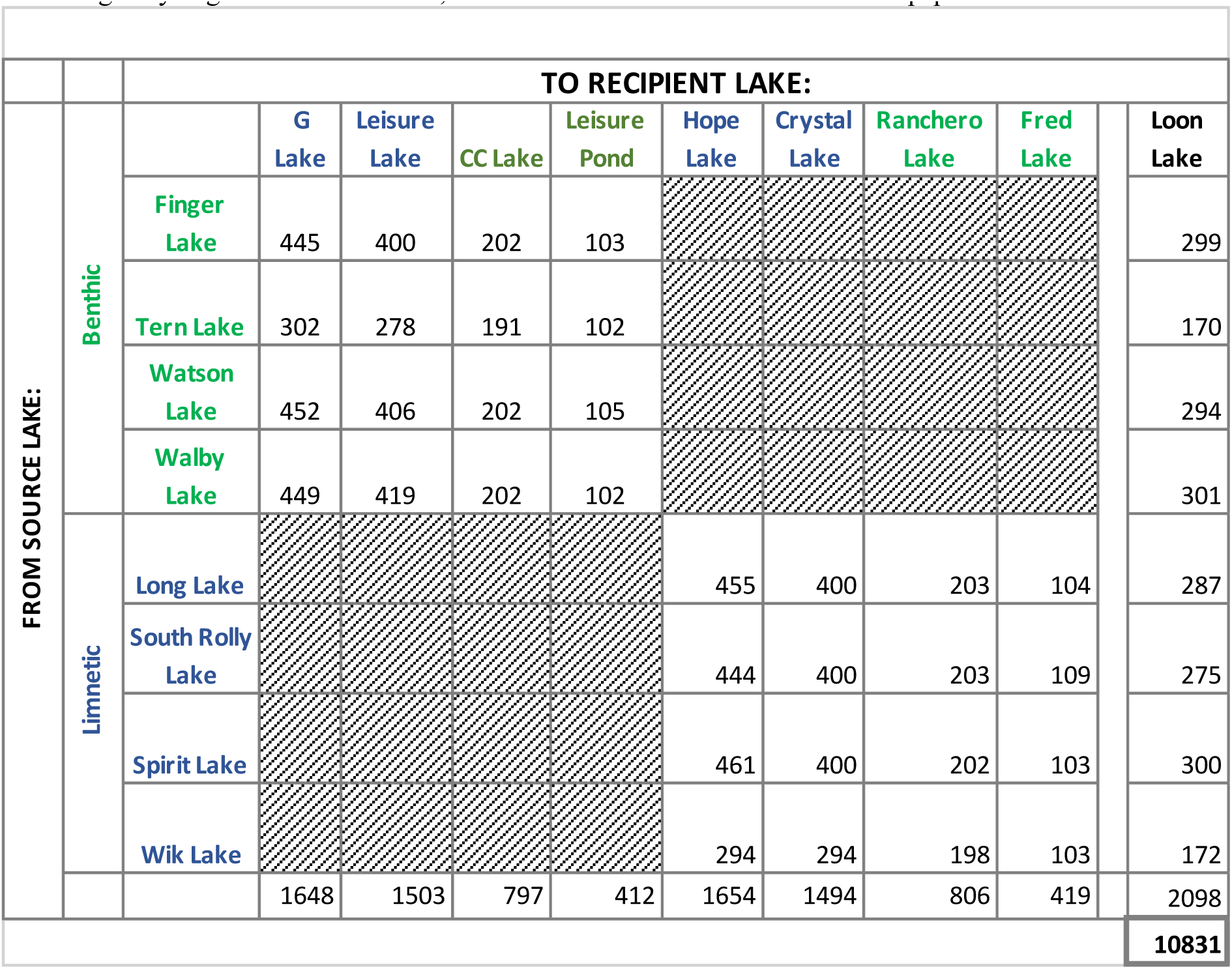
Experimental design showing the numbers of fish transplanted from source to recipient lakes. Source lakes and recipient lakes are color coded by whether each is considered benthic (green) or limnetic (blue). The initial introduction to G Lake failed for unknown reasons, so we repeated the introduction in 2022 and in 2023 confirmed successful establishment, and reintroduced stickleback into G Lake, equally drawn from all source lakes (except Long, where a pike invasion has caused stickleback collapse). Because the second successful G lake population is not chronologically aligned with other lakes, we do not consider G Lake further in this paper.

**Table S2.** Sample sizes of source and recipient lakes through years for fibrosis and infection data. Source Lakes were not sampled in 2020 due to COVID. Recipient lake populations were extinct at the time of sampling in 2019, so were not sampled.

|  |  | **2019** | **2020** | **2021** | **2022** | **2023** |
| --- | --- | --- | --- | --- | --- | --- |
| Recipient Lakes | CC Lake |  | 40 | 100 | 50 | 100 |
|  | Crystal Lake |  | 40 | 100 | 100 | 100 |
|  | Fred Lake |  | 40 | 100 | 50 | 100 |
|  | G Lake * |  | 1 | 0 | 0 | 150 |
|  | Hope Lake |  | 40 | 100 | 50 | 100 |
|  | Leisure Lake |  | 40 | 100 | 100 | 100 |
|  | Leisure Pond |  | 40 | 100 | 100 | 52 |
|  | Loon Lake |  | 40 | 100 | 81 | 100 |
|  | Ranchero Lake |  | 40 | 100 | 50 | 100 |
| Source Lakes | Finger Lake | 100 | 0 | 30 | 30 | 100 |
|  | Jean Lake ** | 97 | 0 | 0 | 0 | 0 |
|  | Long Lake *** | 80 | 0 | 30 | 0 | 0 |
|  | South Rolly Lake | 98 | 0 | 30 | 50 | 100 |
|  | Spirit Lake | 98 | 0 | 100 | 30 | 100 |
|  | Tern Lake | 65 | 0 | 30 | 50 | 65 |
|  | Walby Lake | 100 | 0 | 30 | 30 | 98 |
|  | Watson Lake | 98 | 0 | 30 | 50 | 100 |
|  | Wik Lake | 98 | 0 | 100 | 30 | 77 |

\* The G Lake introduction failed in 2019-2020. A new reintroduction was attempted in 2022 and succeeded. This paper omits G lake from consideration because we lack multiple years of time series data from it.
\*\* Jean Lake was removed from consideration as a source lake after sampling in 2019.
\*\*\* Long Lake stickleback are presumed extinct after pike appeared in 2022 and stickleback catch rates dropped to zero.

**Table S3:** The list of SNPs unique to each source population that were used to infer ancestry of fish from the recipient lakes. The population abbreviations for each lake are: LG (Long), SL (Spirit), SR (South Rolly), WK (Wik), FG (Finger), TL (Tern), WB (Walby), WT (Watson).

| Assay | Population | Chromosome | Position | Unique Base | Allele Frequency |
| --- | --- | --- | --- | --- | --- |
| Limnetic | LG | chrII | 15917728 | T | 0.76 |
| Limnetic | LG | chrIII | 823613 | A | 0.76 |
| Limnetic | LG | chrIII | 9359110 | A | 0.68 |
| Limnetic | LG | chrIV | 20067576 | A | 0.85 |
| Limnetic | LG | chrIV | 13299762 | T | 0.75 |
| Limnetic | LG | chrV | 1753768 | A | 0.66 |
| Limnetic | LG | chrVI | 7405819 | A | 0.64 |
| Limnetic | LG | chrVII | 21415801 | T | 0.69 |
| Limnetic | LG | chrVIII | 14289522 | T | 0.71 |
| Limnetic | LG | chrX | 6603858 | T | 0.68 |
| Limnetic | LG | chrXI | 16193369 | C | 0.91 |
| Limnetic | LG | chrXI | 3081566 | A | 0.80 |
| Limnetic | LG | chrXII | 2375936 | A | 0.72 |
| Limnetic | LG | chrXIII | 8095355 | T | 0.73 |
| Limnetic | LG | chrXIV | 2478199 | A | 0.61 |
| Limnetic | LG | chrXV | 14863116 | T | 0.56 |
| Limnetic | LG | chrXVII | 9235480 | T | 0.61 |
| Limnetic | LG | chrXVIII | 5379465 | T | 0.67 |
| Limnetic | LG | chrXX | 3621731 | C | 0.69 |
| Limnetic | LG | chrXXI | 6464598 | A | 0.58 |
| Limnetic | SL | chrI | 6553442 | T | 0.90 |
| Limnetic | SL | chrII | 7692517 | T | 0.97 |
| Limnetic | SL | chrII | 7692967 | A | 0.96 |
| Limnetic | SL | chrIII | 9231340 | C | 0.93 |
| Limnetic | SL | chrIV | 14839069 | T | 0.95 |
| Limnetic | SL | chrVI | 8431323 | A | 0.95 |
| Limnetic | SL | chrVI | 8425114 | A | 0.94 |
| Limnetic | SL | chrVII | 1817761 | A | 1.00 |
| Limnetic | SL | chrVIII | 3643167 | A | 0.90 |
| Limnetic | SL | chrX | 7342470 | A | 0.78 |
| Limnetic | SL | chrXVI | 7290192 | T | 0.91 |
| Limnetic | SL | chrXI | 931319 | T | 0.98 |
| Limnetic | SL | chrXIII | 591253 | T | 0.90 |
| Limnetic | SL | chrXIV | 11492271 | T | 0.68 |
| Limnetic | SL | chrXV | 504571 | A | 0.80 |
| Limnetic | SL | chrXVI | 10929953 | A | 0.92 |
| Limnetic | SL | chrXVII | 7968281 | T | 0.88 |
| Limnetic | SR | chrI | 26934908 | C | 1.00 |
| Limnetic | SR | chrI | 26952877 | A | 1.00 |
| Limnetic | SR | chrII | 13580477 | T | 0.92 |
| Limnetic | SR | chrIII | 11145910 | A | 0.78 |
| Limnetic | SR | chrIV | 32386335 | A | 0.97 |
| Limnetic | SR | chrIX | 19733496 | A | 0.93 |
| Limnetic | SR | chrV | 9747888 | A | 0.94 |
| Limnetic | SR | chrV | 9747863 | C | 0.94 |
| Limnetic | SR | chrVI | 15534687 | A | 0.79 |
| Limnetic | SR | chrVII | 2508228 | A | 1.00 |
| Limnetic | SR | chrVIII | 18169994 | A | 0.92 |
| Limnetic | SR | chrX | 15470216 | A | 0.84 |
| Limnetic | SR | chrXI | 8508229 | T | 0.96 |
| Limnetic | SR | chrIX | 10856659 | T | 0.93 |
| Limnetic | SR | chrXII | 13556549 | A | 0.95 |
| Limnetic | SR | chrXIII | 3295246 | A | 0.78 |
| Limnetic | SR | chrXIV | 6944009 | T | 0.89 |
| Limnetic | SR | chrXV | 6208432 | T | 0.72 |
| Limnetic | SR | chrXVI | 11846433 | A | 0.84 |
| Limnetic | SR | chrXVII | 6014009 | T | 0.95 |
| Limnetic | SR | chrXVIII | 6861019 | T | 0.77 |
| Limnetic | SR | chrXX | 3592900 | A | 0.99 |
| Limnetic | SR | chrXX | 3813198 | T | 0.99 |
| Limnetic | SR | chrXXI | 11294121 | T | 0.85 |
| Limnetic | WK | chrI | 9646224 | C | 0.88 |
| Limnetic | WK | chrII | 1721718 | A | 0.71 |
| Limnetic | WK | chrIV | 29635767 | T | 0.95 |
| Limnetic | WK | chrIX | 11691955 | A | 0.77 |
| Limnetic | WK | chrV | 9684570 | A | 0.95 |
| Limnetic | WK | chrV | 2195413 | T | 0.88 |
| Limnetic | WK | chrVI | 2510463 | T | 0.82 |
| Limnetic | WK | chrVI | 10480595 | T | 0.80 |
| Limnetic | WK | chrVII | 7026346 | A | 0.88 |
| Limnetic | WK | chrVIII | 6263140 | T | 0.68 |
| Limnetic | WK | chrX | 6995177 | A | 0.75 |
| Limnetic | WK | chrXI | 12958240 | T | 0.65 |
| Limnetic | WK | chrXII | 13741528 | T | 0.61 |
| Limnetic | WK | chrXIII | 14435087 | C | 0.70 |
| Limnetic | WK | chrXIV | 6924894 | T | 0.81 |
| Limnetic | WK | chrXV | 1193653 | A | 0.70 |
| Limnetic | WK | chrXVI | 10258483 | T | 1.00 |
| Limnetic | WK | chrXVII | 7254362 | T | 0.89 |
| Limnetic | WK | chrXVII | 7010766 | A | 0.87 |
| Limnetic | WK | chrXVIII | 14389316 | T | 0.71 |
| Limnetic | WK | chrXX | 13672198 | T | 0.98 |
| Limnetic | WK | chrXX | 13069503 | A | 0.93 |
| Limnetic | WK | chrXXI | 2104869 | C | 0.65 |
| Benthic | FG | chrI | 15704205 | T | 0.66 |
| Benthic | FG | chrII | 8998980 | T | 0.78 |
| Benthic | FG | chrII | 2892706 | C | 0.76 |
| Benthic | FG | chrIII | 11904068 | C | 0.74 |
| Benthic | FG | chrIV | 31662263 | T | 0.65 |
| Benthic | FG | chrIX | 2916678 | T | 0.74 |
| Benthic | FG | chrV | 8194471 | C | 0.77 |
| Benthic | FG | chrVI | 9113425 | T | 0.61 |
| Benthic | FG | chrVII | 20096576 | A | 0.62 |
| Benthic | FG | chrVIII | 7909323 | T | 0.49 |
| Benthic | FG | chrX | 13397132 | A | 0.58 |
| Benthic | FG | chrXII | 2312380 | A | 0.58 |
| Benthic | FG | chrXIII | 3776929 | A | 0.53 |
| Benthic | FG | chrXIV | 13767296 | T | 0.66 |
| Benthic | FG | chrXV | 12722008 | A | 0.93 |
| Benthic | FG | chrXV | 13373000 | A | 0.80 |
| Benthic | FG | chrXVI | 16659604 | C | 0.70 |
| Benthic | FG | chrXVII | 9976657 | A | 0.69 |
| Benthic | FG | chrXVIII | 6306103 | A | 0.85 |
| Benthic | FG | chrXVIII | 14565840 | A | 0.78 |
| Benthic | FG | chrXX | 19635221 | A | 0.86 |
| Benthic | FG | chrXX | 5821398 | A | 0.76 |
| Benthic | FG | chrXXI | 10304559 | A | 0.49 |
| Benthic | TL | chrI | 20114340 | C | 1.00 |
| Benthic | TL | chrI | 20163367 | A | 1.00 |
| Benthic | TL | chrII | 21053847 | T | 1.00 |
| Benthic | TL | chrIV | 5729611 | T | 1.00 |
| Benthic | TL | chrII | 20097809 | A | 1.00 |
| Benthic | TL | chrIX | 348981 | C | 1.00 |
| Benthic | TL | chrV | 2590697 | T | 1.00 |
| Benthic | TL | chrVI | 8817301 | C | 1.00 |
| Benthic | TL | chrVII | 1723789 | A | 1.00 |
| Benthic | TL | chrX | 3733989 | T | 0.98 |
| Benthic | TL | chrXI | 658255 | T | 0.99 |
| Benthic | TL | chrXII | 136786 | A | 1.00 |
| Benthic | TL | chrXIII | 915897 | A | 1.00 |
| Benthic | TL | chrXV | 10269146 | A | 0.99 |
| Benthic | TL | chrXVI | 5926417 | A | 1.00 |
| Benthic | TL | chrXVII | 8487039 | T | 1.00 |
| Benthic | TL | chrIII | 130727 | A | 1.00 |
| Benthic | TL | chrXVIII | 766384 | C | 0.99 |
| Benthic | TL | chrXX | 14862997 | A | 1.00 |
| Benthic | TL | chrXXI | 11630457 | A | 0.99 |
| Benthic | WB | chrI | 11658546 | T | 0.68 |
| Benthic | WB | chrIII | 6828202 | T | 0.67 |
| Benthic | WB | chrIX | 11571874 | A | 0.92 |
| Benthic | WB | chrIX | 11569307 | T | 0.91 |
| Benthic | WB | chrV | 2233732 | T | 0.62 |
| Benthic | WB | chrVI | 11736089 | T | 0.69 |
| Benthic | WB | chrVII | 7165503 | T | 0.80 |
| Benthic | WB | chrVIII | 5789299 | C | 0.63 |
| Benthic | WB | chrX | 651167 | T | 0.63 |
| Benthic | WB | chrXI | 2032558 | A | 0.84 |
| Benthic | WB | chrXIII | 77338 | T | 0.64 |
| Benthic | WB | chrXIV | 5133486 | A | 0.61 |
| Benthic | WB | chrXV | 10053758 | A | 0.63 |
| Benthic | WB | chrXVI | 11188280 | A | 0.93 |
| Benthic | WB | chrXVII | 4145810 | T | 0.67 |
| Benthic | WB | chrXVIII | 4068044 | T | 0.91 |
| Benthic | WB | chrXVIII | 4078056 | T | 0.90 |
| Benthic | WB | chrXXI | 11305884 | T | 0.84 |
| Benthic | WT | chrI | 6268283 | A | 0.74 |
| Benthic | WT | chrIII | 8273044 | A | 0.63 |
| Benthic | WT | chrIV | 25951865 | T | 0.74 |
| Benthic | WT | chrIV | 13840483 | A | 0.72 |
| Benthic | WT | chrV | 7449895 | T | 0.68 |
| Benthic | WT | chrVII | 27191714 | T | 0.66 |
| Benthic | WT | chrVIII | 3764596 | A | 0.79 |
| Benthic | WT | chrX | 3240678 | A | 0.82 |
| Benthic | WT | chrX | 3233056 | A | 0.82 |
| Benthic | WT | chrXI | 11603955 | T | 0.73 |
| Benthic | WT | chrXII | 9432933 | A | 0.71 |
| Benthic | WT | chrXIII | 7970424 | A | 0.70 |
| Benthic | WT | chrXX | 3577020 | A | 0.75 |

**Table S4.** Sample size for lab-based immunization experiments.

| Population | PBS | Alum | NPCGG<br>in Alum |
| --- | --- | --- | --- |
| Finger Lake | 4 | 5 | 5 |
| Long Lake | 9 | 9 | 10 |
| South Rolly Lake | 9 | 8 | 8 |
| Spirit Lake | 11 | 10 | 11 |
| Tern Lake | 9 | 10 | 7 |
| Walby Lake | 11 | 9 | 12 |
| Watson Lake | 11 | 8 | 12 |
| Wik Lake | 6 | 6 | 9 |

**Table S5.** Sample size for lab-based experimental infection assays.

| Population | Experiment | Sample Size | Parasite strain | Fibrosis assayed X days<br>post exposure |
| --- | --- | --- | --- | --- |
| Finger | A | 11 | Finger | 43 |
| Finger | A | 13 | Tern | 42-43 |
| Finger | A | 10 | Walby | 42 |
| Finger | A | 5 | None (copepod only) | 13 |
| South Rolly | A | 7 | Kjerringøy | 44-48 |
| South Rolly | A | 8 | Tern | 43-44 |
| Spirit | A | 6 | Tern | 42-44 |
| Spirit | A | 6 | Walby | 42 |
| Spirit | A | 4 | None (copepod only) | 14 |
| Tern | A | 4 | Kjerringøy | 46-48 |
| Tern | A | 9 | Tern | 43 |
| Tern | A | 8 | Walby | 42-43 |
| Walby | A | 10 | Kjerringøy | 42-48 |
| Walby | A | 12 | Tern | 42-48 |
| Walby | A | 12 | Walby | 42 |
| Walby | A | 5 | None (copepod only) | 15 |
| Watson | A | 6 | Tern | 43 |
| Watson | A | 6 | Walby | 43 |
| Wik | A | 4 | Kjerringøy | 41-43 |
| Wik | A | 6 | Tern | 43 |
| Wik | A | 6 | Walby | 42 |
| Watson | B | 16 | Walby | 30 |
| Finger | B | 7 | Walby | 30 |
| Spirit | B | 15 | Walby | 30 |
| Wik | B | 2 | Walby | 30 |
| Watson | B | 16 | Walby | 30 |
| Finger | B | 7 | Walby | 30 |

